# Co-engagement of TIGIT^+^ immune cells to PD-L1^+^ tumours by a bispecific antibody potentiates T-cell response and tumour control

**DOI:** 10.1101/2023.05.11.540461

**Authors:** Xiaopei Cui, Xi Zhu, Haijia Yu, Jingen Xu, Xiaoyue Wei, Shi Chen, Yangtin Wang, Xiaofang Chen, Yujie Feng, Xiaochen Ren, Liyang Fei, Bin Xie, Mingwei Li, Xue Li, Huifeng Jia, Simin Xie, Li Chen, Yong Cheng, Lei Zhang, Haidong Li, Xiangyang Zhu, Yifan Zhan

**Affiliations:** Drug Discovery, Shanghai Huaota Biopharmaceutical Co. Ltd., Shanghai 201203, China; Peter MacCallum Department of Oncology & Centre for Cancer Research, University of Melbourne, Australia; College of Biology and Pharmacy, Yulin Normal University, Yulin 537000, China

**Author notes:** These authors contribute equally as co-first authors (Xiaopei Cui, Xi Zhu, Haijia Yu, Jingen Xu).

**Keywords:** TIGIT, Bispecfic Antibody, Cancer immunotherapy

## Abstract

Co-targeting PD-1/PD-L1 and TIGIT/CD226 is being pursued to broaden the efficacy of current immunotherapy. Here we demonstrate that a bispecific antibody (BsAb) targeting PD-L1 and TIGIT, HB0036, shows major advantages over the combination of the two parental monoclonal Abs (mAbs). We demonstrated that HB0036 co-engages PD-L1^+^ tumour cells and TIGIT^+^ T cells, and thereby upregulates CD226 on T cells and induces a greater T-cell proliferative response compared to the combination of the parental antibodies *in vitro*. *In vivo*, HB0036 recruits greater amounts of TIGIT antibody in PD-L1^+^ tumours but not in PD-L1^-^ tumours, compared with therapy using the two parental antibodies. We also observed improved tumour control and favourable immunological signatures with HB0036 in syngeneic and xenograft tumour models. Collectively, these findings demonstrate that bispecific antibodies targeting PD-L1 and TIGIT offer superior benefits in cancer immunotherapy compared with therapy using the two parental antibodies. Based on these studies, a phase I clinical trial with HB0036 has been initiated in patients with solid tumours (NCT05417321).

## Highlights

1. BsAb HB0036 enhances the expression of the activating receptor CD226, and substantially increases T cell proliferation and cytokine production when stimulated, compared to T cells exposed to the combination of the two parental mAbs *in vitro*. This advantageous outcome is only evident when tumour cells co-express CD155 and PD-L1, highlighting one of the benefits of BsAb: co-engagement of immune cells and PD-L1^+^ tumour cells.
2. BsAb HB0036 allows greater accumulation of anti-TIGIT Ab at the tumour site through the co-engagement of PD-L1 on tumour cells with the anti-PD-L1 arm of BsAb, leading to high local drug concentrations, when compared to combination therapy with the two parental antibodies.
3. In preclinical tumour models, BsAb HB0036, substantially improves tumour control and is associated with favourable anti-tumour signatures, including reduced infiltration of neutrophils, lower abundance of TIGIT^+^ T cells and increased production of IFN-γ by TILs, when compared to combination therapy with the two parental antibodies.

## Introduction

Avoidance of immune destruction is a hallmark of cancer (1). Overcoming such immune avoidance by checkpoint blockade has revolutionized cancer therapy. For example, inhibitors of either programmed death receptor 1 (PD-1) or its ligand (PD-L1) have shown efficacy against many types of cancer (2,3). A recent study showed that all locally advanced rectal cancer patients with DNA mismatch repair deficiency had strong clinical responses to PD-1 blockade therapy (4). Overall, the objective response rate for PD-1/PD-L1 blockade is only 20-30% in cancer patients (5,6). Combination therapies are therefore being developed to improve response rates (5,6).

Combination therapies include concomitant use of chemotherapy and radiation therapy, cell therapy, agonist biological agents (such as IL-2, IL-15, CD137) or antibodies targeting surface proteins acting as immune checkpoints (CTLA4, TIGIT, TGF-β, LAG3, CD47, CD73 and TIGIT) (7). Recent studies revealed that dual blockade of PD1 and TIGIT enhances T-cell responses and increases inhibition of tumour growth (8,9). Expression and activation of a costimulatory receptor CD226 are mandatory for the effectiveness of such dual blockade of PD1 and TIGIT (8,9). However, despite positive results from multiple pre-clinical studies and early clinical trials, the combination of anti-TIGIT Ab Tiragolumab and anti-PD-L1 Ab Atezolizumab did not offer better progression free survival (PFS), compared to treatmant with Atezolizumab alone in two recent phase 3 lung cancer trials (10). In the face of these setbacks, alternative approaches for targeting TIGIT need to be investigated.

A promising derivative of combination therapy is the use of bispecific antibodies (BsAb), whereby one antibody molecule has two different arms each recognizing a different epitope (11,12). Compared with mAbs, BsAbs can potentially improve specificity and effectiveness, reduce dosages by directly linking immune cells to malignant cells and reduce side effects (12). Nevertheless, there is a lack of solid evidence that BsAbs co-targeting PD-1/PD-L1 and TIGIT/CD226/CD155 pathways can offer better control of tumour burden compared to combinatorial immunotherapy using the two parental mAbs. Therefore, we investigated whether BsAbs targeting PD-L1 and TIGIT could improve immune responses and increase inhibition of tumour expansion compared to treatment with the two parental antibodies. First, we demonstrated that BsAb HB0036 enhanced CD226 expression and proliferation of TCR-stimulated T cells *in vitro* over a great dose range compared to treatment with the two parental antibodies. We then showed that through engagement of its PD-L1 arm, BsAb HB0036 markedly increased the accumulation of anti-TIGIT Ab at PD-L1^+^ tumour sites. Finally, treatment of mice with BsAb HB0036 improved tumour control and was associated with favourable immunological signatures in some preclinical tumour models. Thus, our study demonstrates that BsAb HB0036 engages PD-L1 expressing tumours to TIGIT-expressing immune cells, resulting in the accumulation of anti-TIGIT Ab within the tumour microenvironment, thereby unleashing T and NK cell intratumoural effector functions, consequently leading to better tumour control.

## Results

### Bispecific antibody HB0036 targeting PD-L1 and TIGIT maintains the high affinity and functional activity of parent mAbs

The PD-L1/TIGIT tetravalent BsAb was derived from a humanized anti-PD-L1 antibody described previously (13); NCT04678908) and a humanized anti-TIGIT antibody HB0030 (CTR20212828, patent No 202010387630.4). It was constructed with whole anti-PD-L1 IgG1 and tandem anti-TIGIT scFVs of HB0030 linked to the C-terminus of the heavy chain of the anti-PD-L1 IgG1 in the format of IgG-scFV (**Figure 1A**). The scFv variable kappa region (Vk) and variable heavy region (VH) domains of the anti-TIGIT antibody were connected and tethered to the IgG1 mab by glycine-rich flexible (GGGGS)4 linkers named linker 1. The anti-TIGIT scFv was built in the VL→ VH orientation and joined in-frame via (GGGGS)n linkers named linker 2 where n was 3,4,5. We tested three linker 2s of different length to link the light chain and heavy chain of HB0030 scFVs and found that 900693 had higher purity and better thermodynamic properties (**Table S1**). This reagent was chosen for further study and named HB0036.

**Figure 1.**
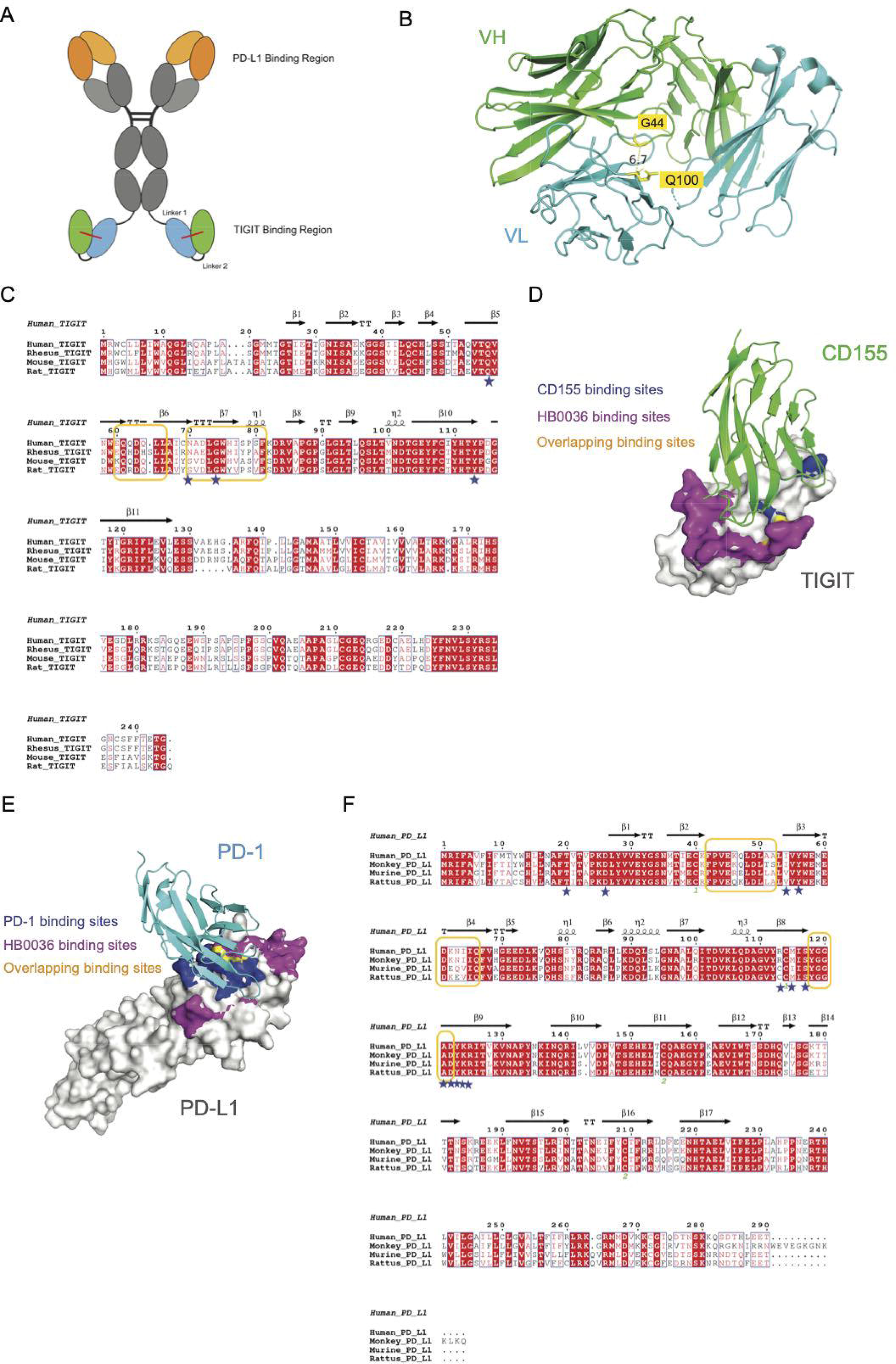
Generation of the tetravalent bispecific antibody HB0036 simultaneously targeting PD-L1 and TIGIT. (**A**) The PD-L1/TIGIT bispecific antibody was constructed by linking a single-chain Fv consisting of heave and light chains of anti TIGIT Ab to the C-terminal domain of the anti-PD-L1 IgG backbone. (**B**) The modeled structure of anti-TIGIT variable region sequences revealed that the distance between Cα atom of VH44 and VL100 was about 6.7 Å, which could contribute to the formation of the disulfide bond between two heave and light chains. (**C**) Sequence conservation and comparison between TIGIT from different species were performed by ClustalOmega program, the secondary structure elements were annotated using ESPript program. Residues of TIGIT involved in binding with HB0036 were marked with yellow rectangular boxes, and residues of TIGIT involved in binding with CD155 were marked with blue stars. (**D**) Epitopes of HB0036 on TIGIT were identified by hydrogen-deuterium exchange (HDX), and the corresponding residues were mapped to the structure of CD155 and TIGIT complex (PDB code: 3UDW) using PyMOL software. (**E**) Epitopes of HB0036 on PD-L1 were identified by hydrogen-deuterium exchange (HDX), and the corresponding residues were mapped to the structure of PD-L1 and PD-1 complex (PDB code: 3BIK) using PyMOL software. (**F**) Sequence conservation and comparison between PD-L1 from different species were performed, the secondary structure elements were annotated. Residues of PD-L1 involved in binding to HB0036 were marked with yellow rectangular boxes, and residues of PD-L1 involved in binding to PD-1 were marked with blue stars.

To increase the stability of HB0030 scFVs, we engineered an inter-chain disulfide bond between the light and heavy chains. It was reported that the distances between the C_α_ atoms for the disulfide bonded cysteines are between 3.0 and 7.5 Å. We used SAbPred tool to build a three-dimensional structure model of the anti-TIGIT Ig variable domain. The predicted distance between C_α_ atom of VH44 and VL100 was about 6.7 Å (14). Using this criterion (distance between C_α_ atom in the range of 3.0 Å and 7.5 Å), a H44-L100 disulfide bond was introduced into the anti-TIGIT scFv (**Figure 1B**) (15). Finally, the epitopes of HB0036 were analyzed by hydrogen deuterium exchange technology. The binding surfaces of HB0036 on TIGIT overlapped with those of CD155, indicating HB0036 could block the binding of CD155 and TIGIT spatially (**Figure 1C&D**). We compared TIGIT sequences of human, Rhesus monkey, mouse and rat and found that sequences of human and Rhesus monkey were more conserved with 88.1% identity, while TIGIT sequences were less conserved between mouse and human, or rat and human (59.0% and 58.0% identity respectively). Similarly, the binding surfaces of HB0036 on PD-L1 overlapped with those of PD-1, and sequences of human and Rhesus monkey were largely conserved with 91.0% identity, while the percentage identity between human and mouse PD-L1 sequences, and the percent identity between human and rat PD-L1 were 69.7% and 70.0%, respectively (**Figure 1E&F**). Thus, HB0036 was deemed likely to block both PD-L1 and CD155-mediated signaling.

As HB0036 bound both PD-L1 and TIGIT, we showed that two targets did not interfere with each other when they interacted with HB0036. When compared with the stoichiometric ratio of samples binding to PD-L1 and TIGIT, no difference was detected in the contact order of the samples exposed to antigens (**Figure 2A&B**). Furthermore, we found that one HB0036 was found to bind to approximately 1.7 PD-L1 monomers and 1.0 TIGIT dimers simultaneously, showing similar binding properties to the parent antibodies against PD-L1 900458 and HB0030 (**Table 1**). Similarly, the valence of antibody binding to antigen was comparable to 900542 (tiragolumab analogue made in house) or Atezolizumab. Next, the binding affinities of HB0036 to TIGIT and PD-L1 were determined and compared to those of the parent anti-PD-L1 mAb and anti-TIGIT mAb, respectively. As for TIGIT, the Kd was determined by Biacore 8k for HB0036, HB0036-PD-L1 complex (after HB0036 incubating with saturated PD-L1 protein), HB0030 and 900542 to be 40.0, 22.8, 13.8 and 65.3 pM, respectively (**Figure 2C**, **Table 2**). As for PD-L1, the KD for HB0036, HB0036-TIGIT complex (after HB0036 incubating with saturated TIGIT protein), 900458 and Atezolizumab were 1980, 2630, 2280 and 1650 pM, respectively **(Figure 2D**, **Table 2)**. Overall, the SPR data indicated that HB0036 had a similar affinity range to PD-L1 and TIGIT when compared to the parent Abs as well as the control Ab 900542 or Atezolizumab.

**Figure 2.**
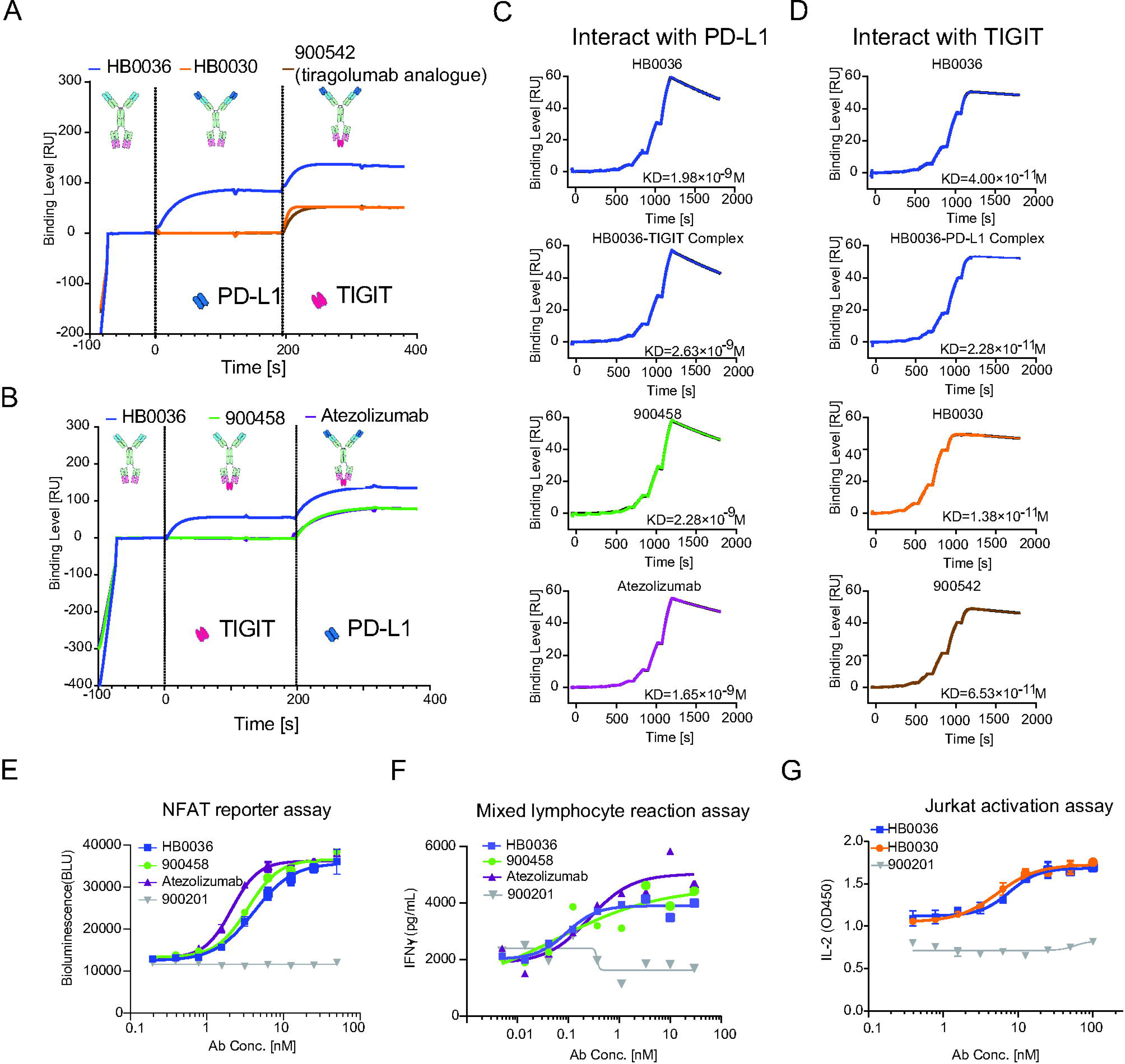
Characterization of binding affinity and functional activity of HB0036. **(A)**. HB0036 (blue) bound in tandem to PD-L1 and TIGIT while HB0030 (yellow) and 900542 (brown) bound TIGIT but not PD-L1. **(B)** HB0036 (blue) bound in tandem to TIGIT and PD-L1 and while 900458 (green) and Atezolizumab (violet) bound to PD-L1 but not to TIGIT. **(C)** kinetics sensorgrams of antibody interacting with human PD-L1. HB0036, HB0036 complex pre-bound TIGIT, 900458 and Atezolizumab were evaluated for PD-L1 binding. **(D)** Kinetics sensorgrams of antibody interacting with human TIGIT. HB0036, HB0036 complex pre-bound human PD-L1, HB0030 and 900542 were evaluated for TIGIT binding. **(E)**. Enhancement of NFAT activation by targeting of PD-L1. CHO-K1-OS8-PD-L1-8D6, expressing human PD-L1 and T-cell activator OKT3 ScFv (OS8) were co-cultured with the Jurkat-NFAT-PD-1-5B8 cells expressing human PD-1 and a luciferase reporter driven by an NFAT-RE in the absence of HB0036, 900458 or Atezolizumab for 6 h. NFAT activation was assessed by determining luciferase activity (bioluminescence) in the supernatant using Bio-Glo^TM^ luciferase assay system. **(F)** Enhancement of IFN-γ production by PD-L1 targeting. CD4^+^ T from one healthy donor were co-incubated with monocyte-derived DC from another healthy donor for 5 days with or without HB0036, 900458, or atezolizumab. IFN-γ in the harvested culture supernatants were evaluated by ELISA. **(H)** Enhancement of IL-2 production by TIGIT targeting. Jurkat-TIGIT-22G8 cells expressing human TIGIT were stimulated with 1 μg/mL PHA-M in the presence or absence of soluble human CD155, with gradient concentrations of HB0036, HB0030 and 900542 for 22 hours. IL-2 in the supernatants was assayed using ELISA. **(G)** ADCC function by NK cells targeting TIGIT. Effector cells NK-92MI-CD16a and target cells (effector: target ratio 3:1) cultured in the presence gradient concentrations of Anti-TIGIT antibodies (HB0036 or HB0030). ADCC activity was measured by using the cytotoxicity LDH Assay Kit.

**Table 1.** Valence of antibody binding to PD-L1 and TIGIT.

| Antibody | Antigen 1 | Antigen 2 | Antigen1:Antibody | Antigen2:Antibody |
| --- | --- | --- | --- | --- |
| HB0036 | PD-L1 | PD-L1 | 1.709 | 0.075 |
|  | TIGIT |  | 1.043 | 1.733 |
|  | PD-L1 | TIGIT | 1.706 | 0.924 |
|  | TIGIT |  | 1.042 | 0.036 |
| 900458 | PD-L1 | PD-L1 | 1.820 | 0.153 |
|  | TIGIT |  | N.B | 1.840 |
| Atezolizumab | PD-L1 | PD-L1 | 1.748 | 0.121 |
|  | TIGIT |  | N.B | 1.740 |
| HB0030 | TIGIT | TIGIT | 1.014 | NA |
|  | PD-L1 |  | N.B | 1.023 |
| 900542 | TIGIT | TIGIT | 0.990 | 0.006 |
|  | PD-L1 |  | N.B | 1.004 |

N.B: no binding

**Table 2.**
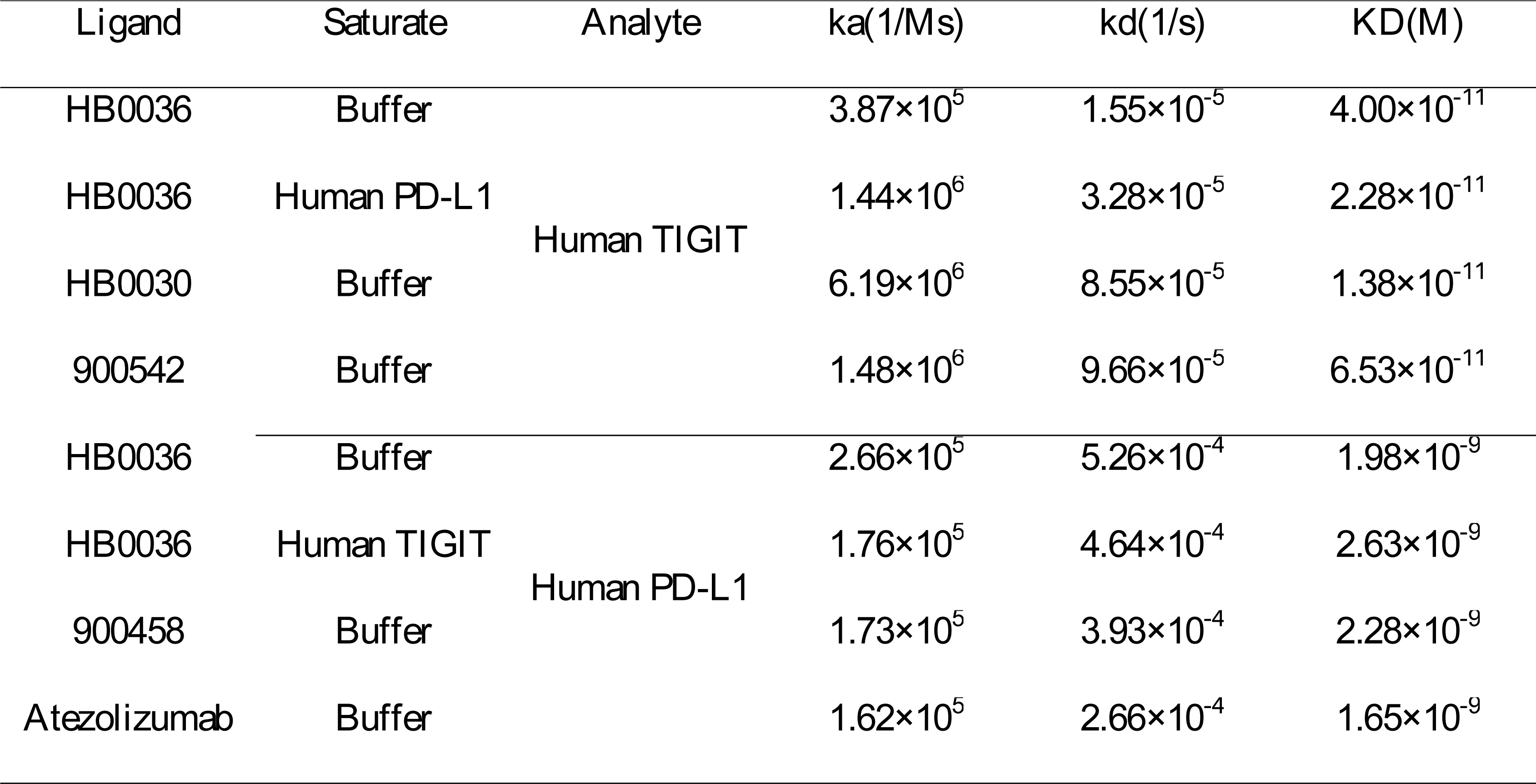
Kinetic results of samples binding to PD-L1 and TIGIT with a 1:1 binding model.

| Ligand | Saturate | Analyte | ka(1/Ms) | kd(1/s) | KD(M) |
| --- | --- | --- | --- | --- | --- |
| HB0036 | Buffer | Human TIGIT | $3.87 \times 10^5$ | $1.55 \times 10^{-5}$ | $4.00 \times 10^{-11}$ |
| HB0036 | Human PD-L1 | | $1.44 \times 10^6$ | $3.28 \times 10^{-5}$ | $2.28 \times 10^{-11}$ |
| HB0030 | Buffer | | $6.19 \times 10^6$ | $8.55 \times 10^{-5}$ | $1.38 \times 10^{-11}$ |
| 900542 | Buffer | | $1.48 \times 10^6$ | $9.66 \times 10^{-5}$ | $6.53 \times 10^{-11}$ |
| HB0036 | Buffer | Human PD-L1 | $2.66 \times 10^5$ | $5.26 \times 10^{-4}$ | $1.98 \times 10^{-9}$ |
| HB0036 | Human TIGIT | | $1.76 \times 10^5$ | $4.64 \times 10^{-4}$ | $2.63 \times 10^{-9}$ |
| 900458 | Buffer | | $1.73 \times 10^5$ | $3.93 \times 10^{-4}$ | $2.28 \times 10^{-9}$ |
| Atezolizumab | Buffer | | $1.62 \times 10^5$ | $2.66 \times 10^{-4}$ | $1.65 \times 10^{-9}$ |

To evaluate the capacity of HB0036 to inhibit the PD-L1 signalling pathway *in vitro*, a luciferase reporter gene system for NAFT activation was employed. CHO-K1-OS8-PD-L1-8D6 cells (expressing human PD-L1 and T-cell activator OKT3 ScFv (OS8)) were used to activate Jurkat-NFAT-PD-1-5B8 cells (expressing human PD-1 and a luciferase reporter driven by an NFAT-RE). NFAT activation was evaluated in the presence or absence of gradient concentrations of HB0036, 900458 and Atezolizumab. HB0036 showed a minor reduction in its potency to reverse the inhibition of T-cell activation (**Figure 2E**). The EC50s of HB0036, 900458 and Atezolizumab were 4.11, 3.43 and 2.10 nM respectively. We also compared HB0036 to 900458 and Atezolizumab in a mixed lymphocyte reaction assay by culturing CD4^+^ T from one healthy donor with monocyte-derived DC from an unrelated healthy donor for 5 days. Production of IFN-γ was similar in all three groups (**Figure 2F**). Together, these data show that HB0036 largely maintained its PD-L1 inhibitory activity. Activity of the anti-TIGIT Ab was evaluated using Jurkat-TIGIT-22G8 cells that over-expressed human TIGIT and could be activated with PHA-M. Jurkat-TIGIT-22G8 cells were incubated with PHA-M and soluble CD155 were measured in the presence of gradient concentrations of HB0036 and HB0030. It was found that enhancement of IL-2 release in the system was similar between HB0036 and HB0030 (**Figure 2G**). These results demonstrate that both the PD-L1 and TIGIT arms of HB0036 are functional, and therefore that HB0036 retained the activities of the two parental mAbs.

### HB0036 is more potent in reversing immune inhibition in T cells and NK cells conferred by CD155 and PD-L1 co-expressing tumour cells *in vitro*

Under TCR stimulation, T cells upregulate PD-1, TIGIT and the related co-activating receptor CD226 and co-inhibitory receptor CD96. Accordingly, when human PBMCs were mitogenically stimulated with antibodies against CD3 and CD28, both CD4^+^ and CD8^+^ T cells showed increased expression of TIGIT, CD226 and CD96 (**Figure 3A**). Similarly, upon TCR stimulation, T cells also upregulated PD-1 (**Figure 3B**). To mimic the tumour microenvironment, we stimulated human PBMCs with antibodies against CD3 and CD28 in the presence of CD155 and PD-L1 expressing tumour cells (**Figure S1A**). We tested the impact of HB0036 or the mixture of equal molar concentrations of the parental mAbs on T-cell proliferation and found that treatment with HB0036 led to a significantly stronger proliferative response in CD4^+^ and CD8^+^ T cells, compared to the combination of anti-PD-L1 and anti-TIGIT mAbs. We also found that the anti-PD-L1 Ab alone but not the anti-TIGIT Ab alone enhanced T cell proliferation (**Figure 3C)**. The differences between HB0036 and the combination of the two parent mAbs were evident over a large range of concentrations (**Figure S1B**). Of note, the maximal enhancement of proliferative response observed in cultures with HB0036 could not be achieved with even high concentrations of 900458 and HB0030 (**Figure S1B**).

**Figure 3.**
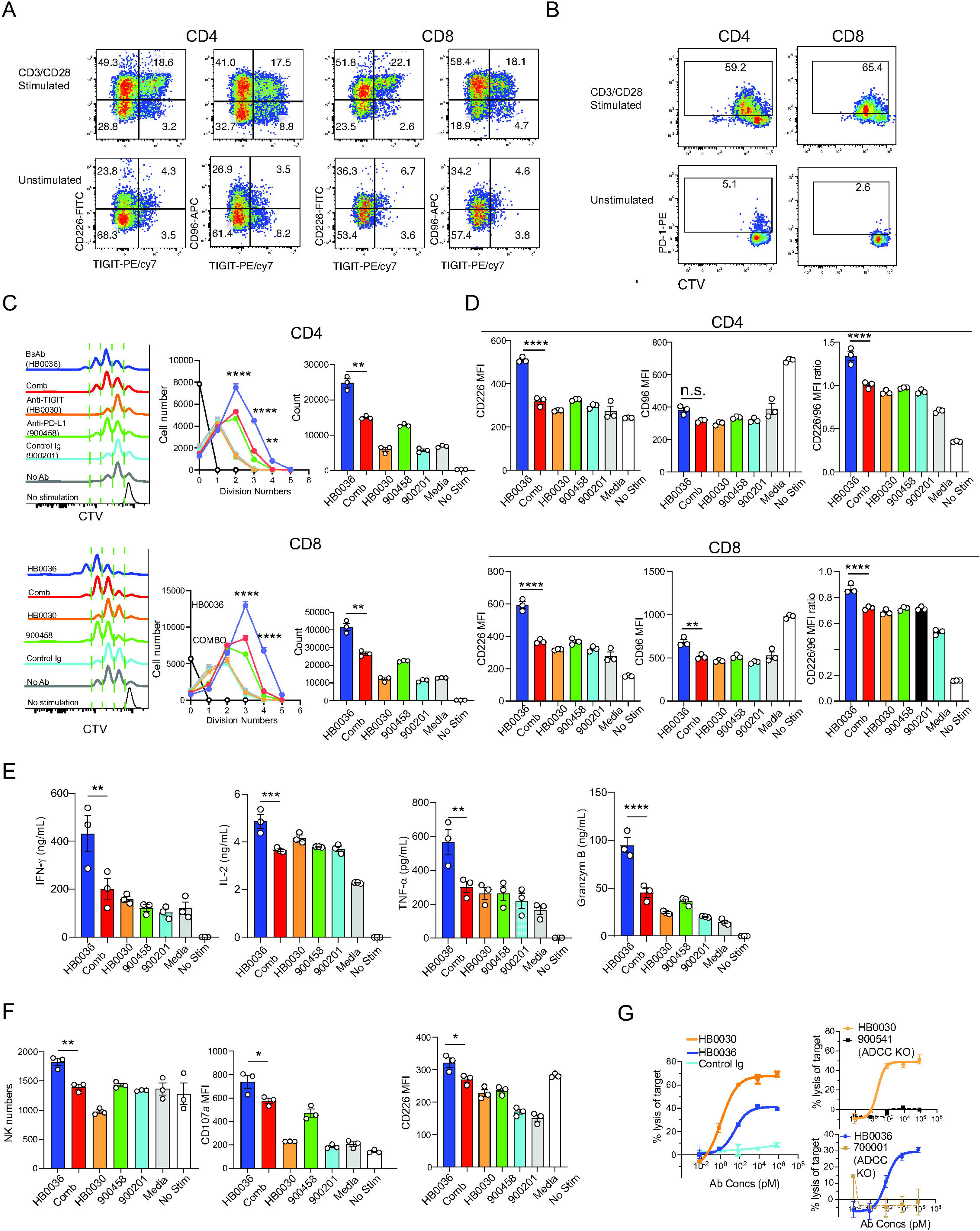
Preferential enhancement of immune responses by bispecific Ab HB0036. (**A&B**) Activation induced expression of co-receptors: Human PBMCs were stimulated *in vitro* with mitogenic antibodies against CD3 and CD28. Harvested cells were stained for surface expression of the indicated markers. FACS plots show expression of CD226, TIGIT, CD96 and PD-1. **(C-F)**. Enhancement of T cell responses *in vitro*: CTV-labelled PBMCs were cultured with mitomycin-treated HT-1080 cells in 96-well plates with indicated Abs at equal molar concentrations. Cells were then stimulated with mitogenic antibodies against CD3 and CD28 for 5 days. Cell proliferation and expression of surface markers were determined. Cytokine production was evaluated by either intracellular staining for cytokines or by using bead arrays. (C). T cell proliferation: histograms display cell division; line graphs show cell numbers at given cell divisions and bar graphs show total recovered proliferating T cells. (D) Bar graphs show the expression (mean MFI+/-SEM) of CD226, CD96 and CD226 MFI/CD96 MFI ratio by CD4^+^ and CD8^+^ T cells. (E) bar graphs show cytokine levels from culture supernatants. (F) Bar graphs show recovered NK cell numbers, expression of CD107a and CD226 by NK cells. *P<0.05, **P<0.01, ***P<0.001, ****P<0.0001 by One-way ANOVA multiple parameter comparison. **(G)** Target cell killing by NK cells. Human NK cells (10^5^/well) as effector cells and Jurkat-TIGIT-22G8 cells (10^5^/well) as target cells were cultured in a 96-well U-bottom plates for 5 h in the presence of serially diluted antibodies as indicated. Lysis of target cells was identified by staining with PI and analyzed by flow cytometry.

The expression pattern of CD155 and PD-L1 by tumour cells was found to predict responsiveness to PD-1 blockade (16). To dissect whether *in vitro* efficacy of HB0036 and combination is also subjected to variation in the expression of CD155 and PD-L1, we used CRISPR/Cas9 system to generate HT1080 without expression of CD155/PD-L1, CD155 only or PD-L1 only by CRISPR/Cas9 (**Figure S1A**). As shown above, when WT HT1080 (CD155^+^PD-L1^+^) was present, HB0036 demonstrated a clear advantage over the combination of the two parent mAbs to reverse immune inhibition (**Figure S1C**). On the other hand, HB0036 did not display significant differences in T-cell proliferation over the combination of the two parent mAbs, even in the presence of a mixture of CD155^+^ and PD-L1^+^ target cells. This indicates that PD-L1 and CD155 need to be expressed on the same cells. In the absence of PD-L1 on tumour cells, CD155 conferred strong immune suppression that could not be reversed by all Abs (**Figure S1C**). In addition, when soluble PD-L1 and CD155 were used to replace CD155 and PD-L1 expressing tumour cells in the culture, suppression of proliferation of CD8^+^ T cells was observed. However, treatment with Abs in all groups resulted in no clear enhancement of T-cell proliferation (**Figure S1D)**. These findings reveal that BsAb promotes greater T cell activation and proliferation in response to tumor cells that co-express CD155 and PD-L1 compared to the combination of the two parent mAbs.

Next, we examined changes of cell phenotype from cultures with WT (CD155 and PD-L1 expressing) tumour cells. We observed that HB0036 but not the combination of the parent mAbs led to higher expression of the co-activating receptor CD226 on both CD4^+^ and CD8^+^ T cells (**Figure 3D & Figure S1E**). Although CD96 was also moderately upregulated in CD8^+^ T cells, the ratio of CD226 to CD96 remained high in T cells treated with HB0036 (**Figure 3D)**. Due to increased numbers of proliferating T cells, the production of cytokines and cytotoxic granule protein including IFN-γ, IL-2, TNF-α and Granzyme B were significantly higher in HB0036 supplemented cultures (**Figure 3E**). The proportions of IFN-γ producing T cells were not significantly different among groups (**Figure S2A-B**). We also observed that HB0036 increased NK cell numbers and increased NK activation based on CD107a expression in PBMC cultures (**Figure 3F**). Moreover, the proportions of IFN-γ producing NK cells was greater in the cultures containing HB0036 (**Figure S2B**). It should be noted that although treatment with anti-TIGIT Ab HB0030 alone had no impact on T cell proliferation, HB0030 enhanced CD107a expression on NK cells (**Figure S2C**) and enhanced NK cell mediated killing of TIGIT expressing target cells in an ADCC-dependent manner (**Figure 3G**). In this aspect, HB0030 was more potent than HB0036 (**Figure 3G**). Overall, HB0036 is more potent in activating T cells when PD-L1 and CD155 are co-expressed on the tumour cells.

### HB0036 enhances intra-tumoural accumulation of anti-TIGIT Ab and reduces intra-tumoural Treg cells

A significant advantage of BsAbs is the ability to bridge immune cells to tumour cells via targeting tumour expressed molecules. PD-L1 is upregulated in the TME on tumour cells and infiltrated leukocytes. To demonstrate the advantage of BsAbs over co-administration of the two parent monoclonal antibodies, hTIGIT/hPD-L1/hPD-1 humanized C57BL/6 mice (n=3 per group) were inoculated *s.c*. with 10^6^ wild type hPD-L1^-^ MC38 cells on the left flank, and 10^6^ hPD-L1^+^ MC38 cells on the right. When the size of the tumour had reached ∼200 mm^3^ (13 days after inoculation), mice were injected intra-peritoneally (i.p.) with either biotinylated HB0036 (683 μg, equal molar concentration to other groups), anti-TIGIT mAb HB0030, anti-PD-L1 mAb 900458, the combination of HB0030 mAb and 900458 mAb, or control Ig 900201 (all 500 μg). Tumours were collected 24 h post Ab injection for analysis of Ab distribution and tumour immune infiltrates (**Figure 4A**). When tumour cells gated as live CD45^-^ cells were analyzed for binding of anti-PD-L1 Ab (detected by PE-streptavidin) and anti-TIGIT Ab (detected by His-tagged TIGIT protein, following APC-anti-His Ab), HB0036 showed abundant binding of both anti-PD-L1 Ab (detected by PE-streptavidin) and anti-TIGIT Ab on hPD-L1^+^ MC38 cells while co-injection of the two parent mAbs only showed strong binding of anti-PD-L1 Ab but not anti-TIGIT Ab (**Figure 4B**). Notably, little Ab binding was detected for all groups on huPD-L1^-^ MC38 tumour cells (**Figure 4B**). In addition, when accumulation of anti-TIGIT Ab at tumour sites was quantified by ELISA, only lysates of hPD-L1^+^ MC38 tumours from mice injected with bispecific HB0036 had significant accumulation of anti-TIGIT Ab (**Figure 4C**). A similar pattern was observed in an *in vitro* system where CHOK1-huPD-L1 cells were incubated with different antibodies (**Figure 4D**).

**Figure 4.**
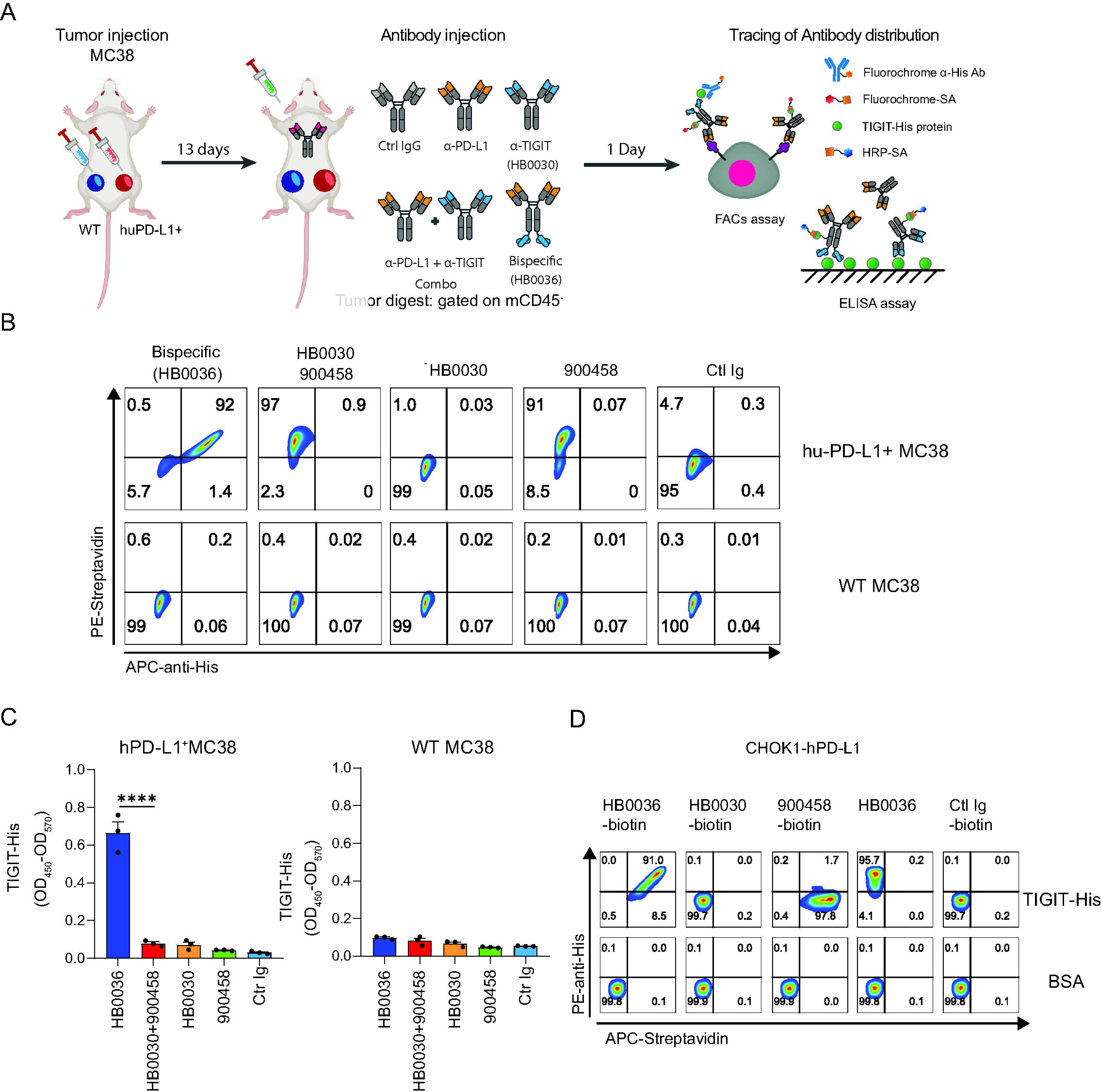
Differences of intra-tumoural accumulation of antibodies between HB0036 and combination of parental antibodies. hTIGIT/hPD-L1/hPD-1 humanized mice were inoculated with hPD-L1^+^ and hPD-L1^-^ WT MC38 cells on the contralateral side for 13 days and were injected with the indicated biotinylated Abs and then harvested after 1 day. Harvested tumours were analyzed for Ab accumulation in the tumour tissue as illustrated in (**A)**. Diagram of experimental design to analyze intratumoural antibody distribution. **(B)**. *In vivo* Ab binding by tumour cells. Tumour digests were incubated with protein TIGIT-His, then stained with PE-Streptavidin (to detect bound biotinylated antibodies), APC anti-His protein (to detect for anti-TIGIT antibodies) and mCD45. FACS plots show Ab binding to tumour cells (gated on live mCD45^-^ cells). (**C**) Accumulation of TIGIT Ab in tumours. Total levels of TIGIT Ab in tumour digests were evaluated by ELISA. **(D)** *In vitro* Ab binding by tumour cells. CHOK1-hPD-L1 was incubated with equal molar amount of biotinylated HB0036, HB0030, 900458 and control Ig, and biotin-free HB0036 respectively for 30 min on ice. Cells were then incubated with human TIGIT protein containing a His tag and incubated on ice for 30 min. Cells were stained with PE streptavidin and APC anti-His tag antibodies on ice for 30 min.

It has been reported that anti-TIGIT Ab reduces the frequency of intra-tumoural T regulatory cells (Tregs) via antibody Fc-dependent cytotoxicity (ADCC) (17). As HB0030 and HB0036 enhanced NK cell mediated killing of TIGIT-expressing cells *in vitro* (**Figure 3G**), we evaluated intra-tumoural Treg cells in the above *in vivo* setting. In the short term (24 h after Ab injection), there were no overt changes in total CD4^+^ T cells **(Figure S3A)**. However, the frequency of Treg cells within intra-tumoural CD4^+^ cells from hPD-L1^+^ MC38 tumours but not from huPD-L1^-^ MC38 tumours was reduced in mice that had been adminitered anti-TIGIT Abs including bispecific Ab, combination of anti-TIGIT and anti-PD-L1 and HB0030 alone (**Figure S3B**). Furthermore, when Treg cells were evaluated for TIGIT expression, hPD-L1^+^ MC38 tumours contained smaller factions of TIGIT^+^ Tregs cell in mice treated with anti-TIGIT Abs while the reduction in TIGIT^+^ Treg cells was less prominent in huPD-L1^-^ MC38 tumours (**Figure S3C**). The reduction in TIGIT^+^ Treg cells was unlikely due to any masking effect of the injected anti-TIGIT Abs as there was no significant binding of His-tagged TIGIT on Treg cells (**Figure S3D)**.

These findings demonstrate that the bispecific HB0036 Ab enhanced intra-tumoural accumulation of anti-TIGIT Ab and reduced the abundance of Treg cells in PD-L1^+^ tumours.

### HB0036 has a stronger anti-tumour effect in a murine colorectal carcinoma model compared to treatment with the parent mAbs

We have shown above that PD-L1 targeting via HB0036 resulted in greater intra-tumoural accumulation of anti-TIGIT Ab and that HB0036 was more potent at reversing immune inhibition exerted by PD-L1/CD155^+^ tumour cells. We next evaluated whether HB0036 could be better at controlling tumour growth than the combination of the parent mAbs. To this end, we firstly investigated the effects of anti-TIGIT Ab HB0030, anti-PD-L1 Ab 900458, as well as combination of HB0030 with anti-PD1 Ab or anti-PD-L1 Ab on tumour growth. Using a subcutaneous model of the murine colorectal carcinoma cell line CT26 in hTIGIT/hPD-1 BALB/c mice, we found that HB0030 with ADCC function caused potent inhibition of tumour growth compared to its ADCC-incompetent counterpart (**Figure S4A**). In the subcutaneously implanted CT26-hPD-L1 tumours in hPD-1/hPD-L1 BALB/c mice, anti-PD-L1 Ab 900458 also inhibited tumour growth (**Figure S4B**). When HB0030 was co-injected with anti-PD-1 Ab Keytruda in the CT26 tumour model (**Figure S4C**) or anti-PD-L1 900458 was used in the MC38-hPD-L1 model in hPD-1/hPD-L1/hTIGIT humanized C57BL/6 mice (**Figure S4D**), we observed enhanced inhibition of tumour expansion (as reflected by reduced tumour volume and tumour weight) compared to treatment with HB0030 alone.

We then compared side by side inhibition of tumour growth by HB0036 vs the combination of the two parent Abs. Inhibition of tumour expansion was first evaluated in a syngeneic mouse model with hTIGIT/hPD-1/hPD-L1 humanization using the syngeneic carcinoma hPD-L1 CT26 cells. We observed significantly reduced tumour volume and tumour weights in the HB0036 treatment group (**Figure 5A**). To dissect the immunological parameters correlating with inhibition of tumour expansion, we examined tumour infiltrating leukocytes (TILs) in control, HB0036 treated and mAb combination treated mice. First, we found that the numbers of TILs (calibrated to the same tumour weight) were greater in the HB0036 and mAb combination treatment groups, compared to the control group (**Figure 5B, Figure S5A**). Furthermore, the numbers of TILs correlated negatively with tumour weight (**Figure 5B**), indicating that the abundance of TILs is critical to inhibit tumour expansion.

**Figure 5.**
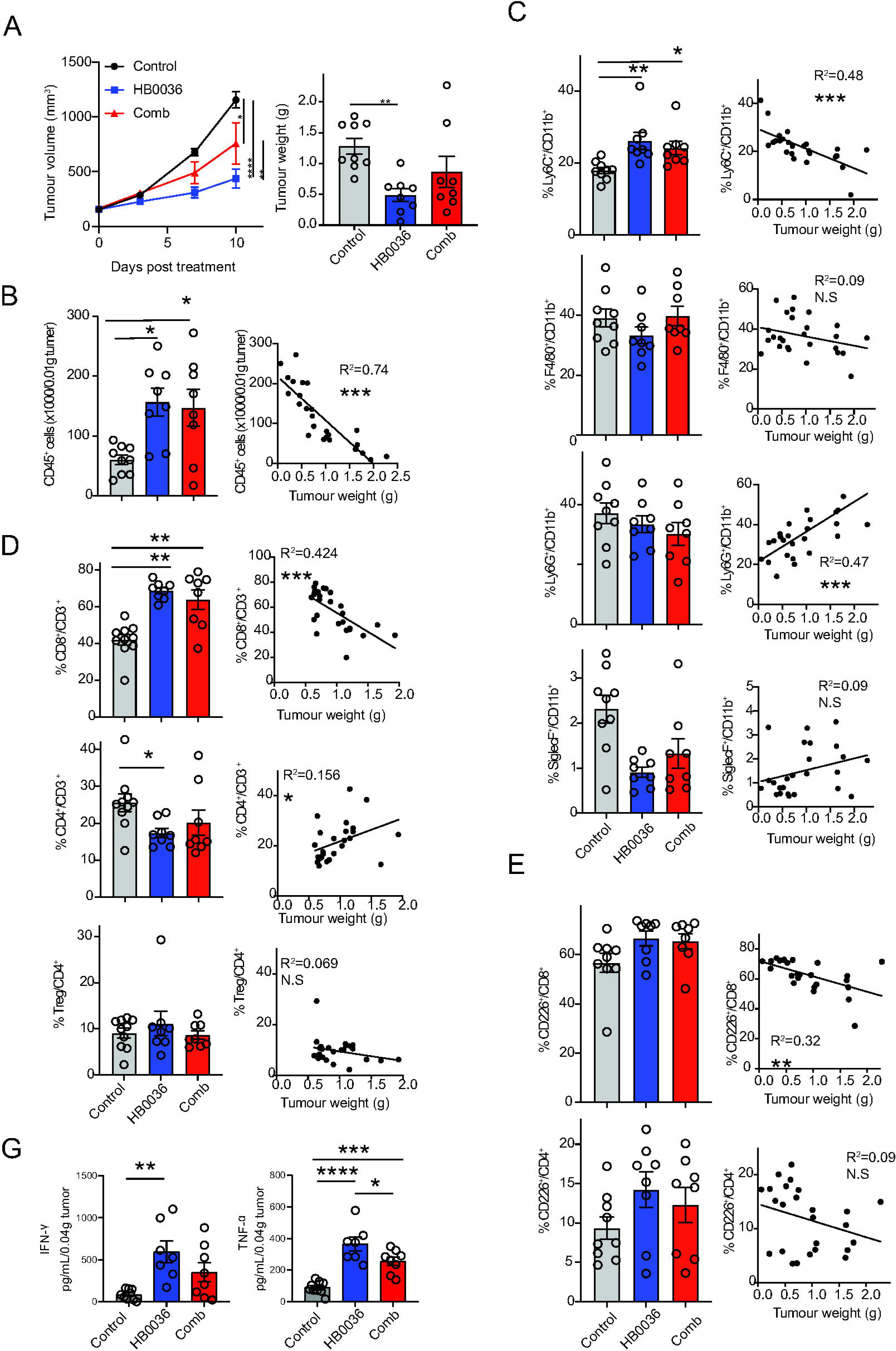
Differences in inhibition of tumour expansion and TILs between HB0036 and combination in the CT26-hPD-L1 syngeneic carcinoma model. BALB/c-hPD1/hPDL1/hTIGIT mice were inoculated with 1×10^6^ CT26-hPDL1 cells on the flank. Treatment (n=8-9) was started when tumour sizes were an average TV of 160 mm^3^. Equimolar dose of Abs (13.7 mg/kg body weight HB0036 or 10 mg/kg 900458 and 10 mg/kg body weight HB0030, twice a week, total 4 doses) were given intraperitoneally. At the end of the experiment, tumour weights were recorded. Tumours were digested for TIL analysis. **A**.Ttumour volumes and tumour weights. P values for tumour volumes were calculated with Prism two-way ANOVA *Tukey’s multiple-comparisons test*. P values for tumour weights were generated with one-way ANOVA multiple *multiple-comparisons test*. *P<0.05, **P<0.01, ***P<0.001, ****P<0.0001. **B-E**. The abundance of TILs [left] and the relationship to tumour weights [right]. B. The numbers of CD45^+^ TILs calibrated to the same tumour weight shown in bar graphs. Linear regression was used to model the relationship between the numbers of CD45^+^ TILs and tumour weights. C. The relative abundance of myeloid cell subsets and its relationship to tumour weights. D. The relative abundances of T cell subsets and its relationship to tumour weights. E. The relative abundance of CD226-expressing T cell subsets and their relationship to tumour weights. P values for TIL abundance were generated with one-way ANOVA *multiple-comparisons test*. Linear aggression was used to generate R^2^ and P values between two parameters. *P<0.05, **P<0.01, ***P<0.001.

Next, we analyzed the relative abundance of different immune cell subsets amongst CD45^+^ TILs. Within the myeloid compartment, Ly6C^+^ monocytes were more abundant in the treated mice but there were no differences between the two different treatment groups **(Figure 5C, Figure S5B)**. The other two major CD11b^+^ subsets, F4/80^+^ macrophages and Ly6G^+^ granulocytes, were not different from the control group **(Figure 5C, Figure S5B)**. A minor population of SiglecF^+^CD11b^+^ myeloid cells were present in TILs. The treated groups contained fewer of them than the control group. Notably, the abundance of Ly6C^+^ monocytes among 4 populations of myeloid cells showed a negative correlation to tumour weight, while Ly6G^+^ granulocytes showed a positive correlation to tumour weight **(Figure 5C, Figure S5B)**. Within the T cell compartment, the two treated groups contained higher proportions of CD8^+^ T cells while the HB0036 treated group contained lower proportions of CD4^+^ T cells than the control group (**Figure 5D; Figure S5C**). Amongst the CD4^+^ T cells, Foxp3^+^ Treg cells were comparable amongst the three treatment groups (**Figure 5D; Figure S5C**).

As CD226 is critical for the anti-tumour effect of both TIGIT and PD-1 inhibitors (9), we evaluated CD226 expression by T cells. A high proportion of CD8^+^ T cells expressed CD226 whilst CD226 expression was only found in 10-15% of CD4 T cells (Figure 5E; **Figure S5D**). The percentages of CD226^+^ subsets trended higher in the treated groups compared to the control group; however, none of the differences reached statistical significance (**Figure 5E**). Nevertheless, CD226 expression by CD8^+^ T cells was negatively correlated with tumour weight (**Figure 5D**). Further characterization of the TILs revealed that the frequency of CD49b^+^ NK cells was not impacted in the different groups (**Figure S5E**). Unexpectedly, our data suggest that NK abundance is not a favourable indication of inhibition of tumour expansion (**Figure S5E**). Finally, the DC subsets (XCR1^+^ cDC1 and SiglecH^+^ pDCs) were the least abundant immune cells present amongst the TILs in all groups (**Figure S5F**).

In addition to phenotypic analysis, tumours were digested and cultured in the presence of TLR agonists to evaluate immunological function of TILs. We found that TILs from the treated groups were able to produce significantly higher levels of IFN-γ and TNF-α (**Figure 5G**). Overall, the treated groups showed favorable immunological signatures, particularly with higher abundance of TILS. Despite the fact that the sub-cellular composition of TILs from mice treated with BsAb or the mAb combinatorial therapy showed only minimal differences, we were able to show that effector cytokines were significantly increased in mice that have been treated with BsAb compared to mice that received the two parent mAbs.

### HB0036 has a stronger anti-tumour effect than mAb combination therapy in the BxPC-3 xenograft model

We next compared tumour control by HB0036 versus the combination of the two parent mAbs in a subcutaneous BxPC-3 xenograft model in human PBMC humanized *Prkdc^scid^Il2rg^null^* (NPG) mice. In this model, we observed that both HB0036 and the combination of the two parent mAbs resulted in smaller tumour volumes, compared to the groups treated with control Ig. Interestingly, HB0036 exerted stronger inhibition of tumour expansion than the combination of the two parent mAbs, although the differences were modest (**Figure 6A**). To evaluate immunological changes that could be associated with differences in the extent of inhibition of tumour growth, we analyzed TILs. TILs contained approximately 2% for mouse myeloid cells and 10% of human lymphocytes in all three groups (**Figure 6B, C&E**). Mouse CD11b^+^ myeloid cells could be divided into two main subpopulations, including Ly6G^+^ neutrophils and Ly6G^-^ cells and Ly6G^-^ cells, which contained Ly6C^+^ monocytes and F4/80^+^ macrophages (**Figure 6B&D**). HB0036 treated mice contained a smaller fraction of Ly6G^+^ neutrophils than the control group and higher percentages of CD11b^+^Ly6G^-^ cells (**Figure 6B&D**). The proportions of human T cells, including both CD8^+^ and CD4^+^ T cells amongst TILs, were not different between the different groups (**Figure 6E&D**). A notable difference was seen in the expression of TIGIT within T cell subsets. For both CD8^+^ and CD4^+^ T cells, the proportions of TIGIT^+^ cells in the HB0036 treated group were significantly decreased compared to the control group or the group treated with the combination of the two parent mAbs (**Figure 6F&G**). In contrast to the observations made above on mouse CD4 T cells (**Figure 5E**), we found that T cells from PBMC-humanized mice expressed high levels of CD226. However, expression of CD226 did not differ between groups (**Figure 6F&G, Figure S6A**). CD96 was expressed on 20-30% of the human T cells but no differences were found between the treatment groups (**Figure 6F&G, Figure S6A**). Improved inhibition of tumour expansion was also observed in xenografts of the human bladder carcinoma cell line HT1367 in PBMC humanized mice (**Figure S6B**).

**Figure 6.**
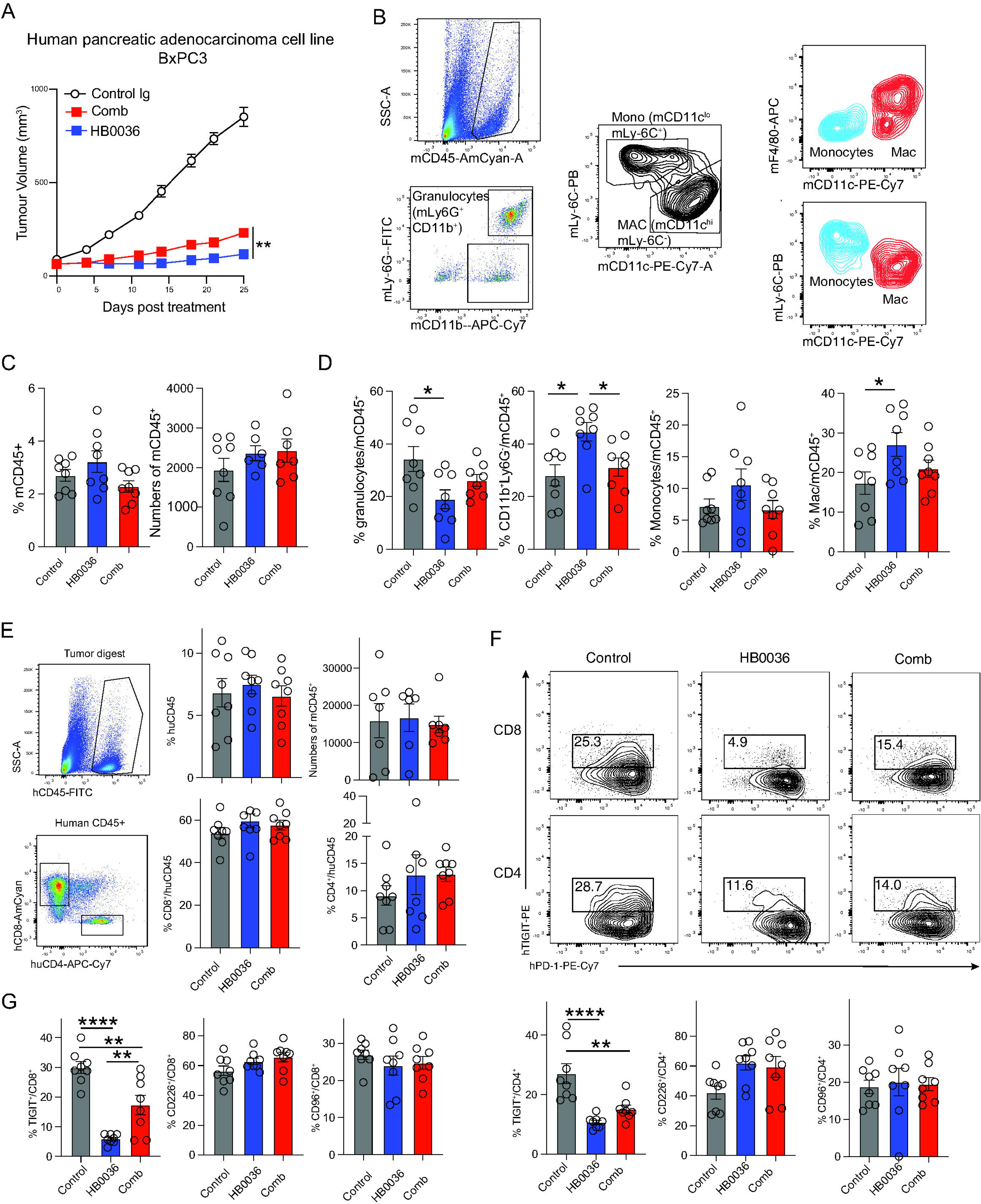
Differences in inhibition of tumour expansion and TILs between HB0036 and Ab combination treatment in the BxPC-3 xenograft model in PBMC humanized mice. BxPC-3 (1×10^7^) in Matrigel were inoculated subcutaneously into NPG mice on the right flank. When tumours had grown to an average size of 80 mm^3^, tumour-bearing mice were randomly divided into 4 groups (n=8 each) and injected i.v with 1×10^7^ PBMCs. Treatments started on the same day and antibodies were dosed twice weekly for 4 weeks intraperitoneally. A. Tumour volume. **P<0.01 by two-way ANOVA Tukey’s multiple-comparisons test. B-D. Analysis of TILs of mouse origin. B. FACS plots show the gating of mouse myeloid cells within mouse CD45^+^ TILs. C. The percentages and the numbers of mouse TILs, D. The percentages of mouse CD11b^+^ myeloid cell subsets. *P<0.05 by one-way ANOVA multiple T test. E-G Analysis of TILs of human origin. E. FACS plots show gating of T cell subsets. Bar graphs show the percentages and/or the numbers of human CD45^+^ TILs and T cell subsets. F. FACS plots show the expression of TIGIT and PD-1 by human CD8^+^ and CD4^+^ T cells. G. Bar graphs show the percentages of human T-cell subsets with expression of TIGIT, CD226 or CD96. **P<0.01, ****P<0.0001 by one-way ANOVA multiple-comparisons test.

Collectively, our investigations using both mouse and human tumour models revealed some differences in the composition of immune cells associated with functional differences between HB0036 treated mice and the mice treated with the combination of the two parent mAbs. These differences included reduced infiltration of neutrophils, lower numbers of TIGIT^+^ T cells and increased production of IFN-γ by TILs. Some or all of these attributes could contribute to the better tumour control by HB0036.

### HB0036 has reduced half-life in mice and cynomolgus monkeys

A factor independent from the mode of action but influencing the outcome of tumour control is the half-life of an Ab and the development of anti-drug antibodies (ADA). Particularly, in this study we tested the efficacy of humanized Abs in pre-clinical murine tumour models. In BALB/c mice, we found that HB0036 had a shorter half-life compared to HB0030 and the anti-PD-L1 Ab 900458 at certain concentrations when given equal molar amounts of Abs (**Figure S6C**). Furthermore, mice treated with HB0036 at lower doses had detectable ADA at 21 days after injection (**Figure S6D**). However, this is likely due to mouse anti-human IgG responses, so ADA are less likely to occur in humans treated with human Abs. To move towards clinical applications, we also performed pharmacokinetic-pharmacodynamic (PKPD) analysis in non-human primate cynomolgus monkeys. When given equal amounts (3 or 10 mg/kg) as a single injection, the concentrations of circulating Abs were comparable between anti-TIGIT mAB HB0030 and BsAb HB0036 over the course of 6 days. Subsequently, the concentrations of HB0036 dropped faster than those of HB0030 (**Figure S6E**). In experiments with multiple Ab infusions, the HB0036 group with lower dose (10 mg/kg) had lower recovery compared to the HB0030 group with the same dose at 2 h after the last injection (day 29). This may indicate the development of ADA following the treatment of non-human primates with HB0036. Overall, these preclinical data indicate that BsAb may have a shorter half-life and that their application may result in the development of ADA. It remains to be addressed whether tailored dosing regimens could circumvent these potential caveats.

### Human tumours exhibit varied pattern of expression of CD155 and PD-L1

Our data highlight that the efficacy of HB0036 or the combination therapy with the two parent mAbs to unleash T cell responses is dictated by the expression of CD155 and PD-L1 by tumour cells. It has been shown that the expression pattern of CD155 and PD-L1 in non-small cell lung carcinoma (NSCLC) tumours predicts response to PD-1 blockade, with patients with tumours expressing PD-L1 without CD155 more likely to respond (16). As for both mAb combination therapy and bispecific Ab therapy, identification of patient cohorts who can benefit from therapy should be pursued. To this end, we analyzed expression of mRNAs encoding CD155 and PD-L1 in patient derived xenografts and human cancer cell lines from the data base of CROWN BIOSCIENCE (https://www.crownbio.com/databases) and CCLE (https://depmap.org/portal/). Expression of PD-L1 was very variable across the different PDX cancers with PDXs of human lymphoma and lung cancers having the highest PD-L1 expression (**Figure 7A**). Similar patterns were observed across a range of human cancer cell lines (**Figure 7B**). On the other hand, expression of CD155 was universally high in solid tumours but low in blood tumours (**Figure 7A&B**). Analysis of co-expression revealed that lung cancers contained a high proportion of PD-L1^hi^ and CD155^hi^ cells while pancreatic, liver and colorectal cancers showed a dominant PD-L1^lo^ and CD155^hi^ cohort (**Figure 7C&Figure S7A**). We selected several human cancer lines to analyze co-expression of PD-L1 and CD155 and demonstrated variable patterns of expression (**Figure 7D**). To complicate this further, the CD155-TIGIT signaling axis consists of many other members that positively and negatively regulate T cell activation. Expression of these molecules also varies greatly among different tumour types (**Figure S7B&C**). As expression of PD-L1 and CD155 has been shown to be associated with treatment efficacy (16), it could be that expression of these two markers as well as related molecules contributes to variation in treatment benefit.

**Figure 7.**
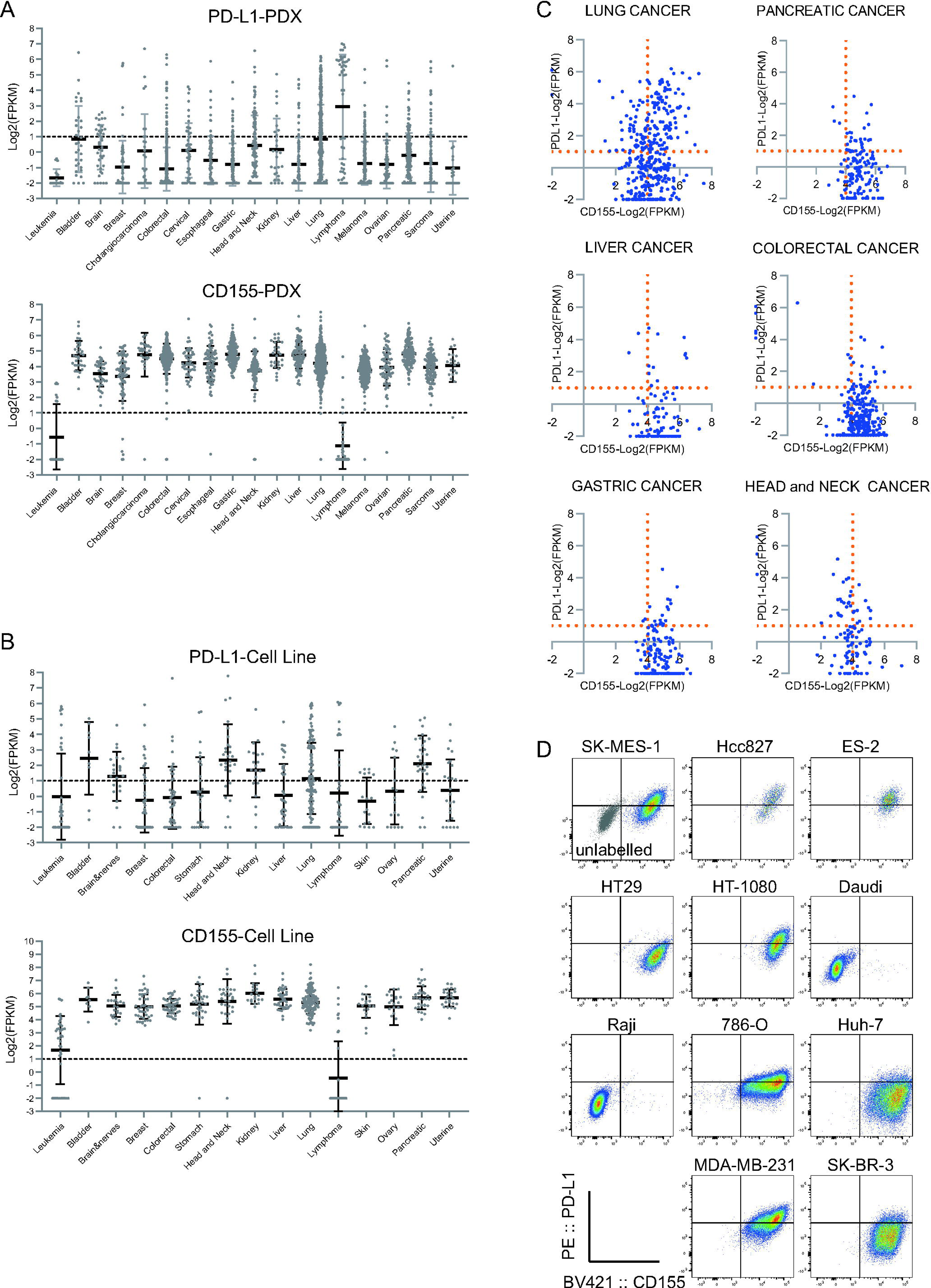
Expression of TIGIT ligand CD155 and PD-L1 on tumour cells. **A-C**. PD-L1 and CD155 mRNA expression of cancer PDX and cancer lines (https://hubase.crownbio.com/GeneExpressionRNAseq/Index) were analyzed. **(A)** PDX data, **(B)** cell line data (https://xenobase.crownbio.com/GeneExpressionRNAseq/Index), **(C)** co-expression pattern of PD-L1 and CD155 of PDX samples. (**D**). Selected human cancer cell lines were analyzed for surface expression of PD-L1 and CD155.

## Discussion

Here we explored the anti-tumour effects of the TIGIT and PD-L1 targeting bispecific Ab HB0036 comparing it to combination therapy using the two parental antibodies. *In vitro*, we demonstrate that BsAb HB0036 increased T cell proliferation in the presence of PD-L1-and CD155-expressing tumour cells compared to mAb combination therapy*. In vivo*, we show that the anti-PD-L1 arm of the bispecific Ab increases the availability of anti-TIGIT at the PD-L1^+^ tumour site and offers improved tumour control associated with favourable immunological changes. This approach may provide a strategy to improve the efficacy of dual blockade of PD-1/PD-L1 and TIGIT/CD155 pathways in cancer immunotherapy.

Most solid tumours express high levels of CD155 and some express PD-L1 (**Figure 7A&B**). To mimic the tumour microenvironment, we included CD155 and PD-L1 expressing HT1080 tumour cells in cultures containing human PBMCs. In this setting, T cell proliferation in response to TCR stimulation was significantly higher in cultures with HB0036 compared to cultures containing the two parent mAbs. Somewhat unexpectedly, when CD155 and PD-L1 were not expressed on the same tumour cells (i.e. CD155^+^PD-L1^-^ and CD155^-^PDL1^+^ HT1080 cells), the advantage of HB0036 treatment was abrogated. This implies that co-engagement of TIGIT on T cells by the anti-TIGIT arm of the BsAb and PD-L1 on tumours via the anti-PD-L1 arm of the BsAb is preferred for maximal immune cell stimulation, probably by facilitating the interaction between CD226 on T cells and CD155 on the tumour cells. In the absence of CD155 on tumour cells, blocking PD-L1 enhanced T-cell proliferation but HB0036 had no advantage over mAb combination treatment. In the absence of PD-L1, both HB0036 and mAb combination treatment had little ability to enhance T-cell proliferation. Overall, the therapeutic value of dual blocking of TIGIT and PD-L1 might depend on the expression pattern of CD155 and PD-L1 on tumour cells. HB0036, via co-engagement of T cells and tumour cells may have an advantage over the mAb combination in patient cohorts with high expression of both CD155 and PD-L1 on tumour cells.

We also noted that enhanced T cell proliferation caused by treatment with HB0036 *in vitro* was associated with higher expression of the costimulatory receptor CD226. As expression and activation of CD226 is required for induction of CD8^+^ T cell response (8, Banta, 2022 #1957), it could be therefore expected that T cells with elevated expression of CD226 in culture with HB0036 had a stronger proliferative response in response to TCR stimulation. CD226 phosphorylation has been shown to be influenced by competition of CD155 binding by the extracellular domain of TIGIT (9). In our case, co-engagement of TIGIT by HB0036 could allow CD155 interacting with CD226 to deliver positive signalling for T cell activation. In support of this idea, in PBMC cultures with soluble CD155 and PD-L1, HB0036 showed no advantage of increasing CD226 and T cell proliferation. Notably, immune suppression enforced by soluble CD155 and PD-L1 was relatively weak compared to that by CD155 and PD-L1 present on the plasma membrane of tumour cells. Of note, it has been reported that CD226 expression can be regulated by at least two different mechanisms (18). Transcriptionally, Eomes is able to directly interact with regulatory elements of the gene encoding CD226 (18,19). Accordingly, Eomes overexpression caused an increase in the percentages of CD226^−^ CD8^+^ T cells (18). Post-transcriptionally, CD155 induces ubiquitination of CD226 via CBL-B and thereby primes it for proteasomal degradation (19). Currently, we have not explored how BsAb HB0036 might increase CD226 expression on activated T cells.

We demonstrated that treatment HB0036 *in vivo* resulted in greater accumulation of TIGIT Ab in the form of bispecific Abs at PD-L1^+^ tumour sites compared to combination treatment with the parent mAbs. Accordingly, we observed improved inhibition of tumour expansion in groups receiving HB0036 in both syngeneic and xenograft tumour models. We further looked into immunological parameters in effector memory T cells (Tem) that may be associated with inhibition of tumour expansion. In the syngeneic CT26-hPD-L1 tumour model, when tumour cell suspensions (from equal weight of tumour) were cultured, the production of IFN-γ and TNF-α from the treated group was significantly higher than from the control group while production of TNF-α was higher in the HB0036 group compared to the mAb combination group. At the cellular level, the treated groups contained more TILs and higher proportions of CD8^+^ T cells. These two parameters favour tumour control.

Conversely, CD49^+^ NK cells and Foxp3^+^ Treg cells, two cell populations known to be targeted by TIGIT blockade, were less affected by treatment with HB0036 or the mAb combination. Interestingly, representation of Ly6C^+^ monocytes within myeloid cell compartment was higher in the treated groups. In the BxPC-3 xenograft tumour model with PBMC humanization, TILs were dominated by human lymphocytes with very few Treg cells and NK cells. Within the T cell compartment, a notable difference found is that the HB0036 group contained lower percentages of TIGIT^+^ cells. Considering, that the expression of TIGIT and PD-1 is a characteristic of exhausted CD8^+^ T cells and is often found in TEM (9,20), reduced TIGIT expression could shift the balance of CD226/TIGIT to immune cell activation. The myeloid cell compartment of TILs in the tumour xenograft models mainly consisted of mouse cells. The HB0036 treated group contained lower percentages of Ly6G^+^ granulocytes. It is not clear how human T cells impacts the phenotype of mouse myeloid cells in the TEM.

Our *in vitro* data showed that blockade of TIGIT alone has limited impact on T cell proliferation. This could be due to the lack of targeted immune cells in the system as TIGIT had been shown to require the presence of dendritic cells to affect T-cell responses (21), although agonistic anti-TIGIT Ab had been found to suppress T-cell response independent of APCs (22). Nevertheless, the anti-TIGIT antibody in our system did induce ADCC-mediated killing of TIGIT expressing tumour cells. *In vivo*, blockade of TIGIT alone had variable efficacy in the inhibition of tumour expansion. In the CT26 tumour model, blockade of TIGIT alone as well as blockade of PD-1/PD-L1with anti-PD-L1 antibody alone only marginally inhibited tumour growth (8). On the other hand, we did observe better tumour control with treatment of anti-TIGIT Ab HB0030 in a similar model, depending on intact Fc function. Depletion of intra-tumoural Treg cells by a Fc-competent anti-TIGIT Ab is recognized as a mechanism that contributes to the inhibition of tumour expansion exerted by the anti-TIGIT Ab (17). In our *in vivo* experiments using h-PD-L1 MC38 tumour cells, we also observed a selective and acute reduction in intra-tumoural Treg cells, particularly the TIGIT^+^ ones, upon administration of a single dose of either the two mAb or BsAbs. However, in our experiments with hPD-L1 CT26 tumour cells, there was no clear difference in Treg cell representation within the TILs when tumours were harvested after treatment with multiple doses of Ab. A question remains whether anti-TIGIT Abs with active FcγR could also kill TIGIT^+^ effector cells. It could be that tumour control by anti-TIGIT Abs with active FcγR depends on the net outcome of Treg cell depletion vs the killing of effector T cells. Of note, several anti-TIGIT Abs with inactive FcγR have also entered into clinical trials (23). Notably, TIGIT is associated with NK cell exhaustion in tumor setting. Blockade of TIGIT resulted in potent tumor-specific T cell immunity in an NK cell-dependent manner (24). Given that PBMC humanized mice lack reconstitution of human NK cells, the anti-tumor role of TIGIT blockade, either in the form of bispecific or combination of parental mAbs, may not be adequately fulfilled in PBMC humanized mice with tumour xenografts.

TIGIT functions in a complex context with the co-existence of related co-inhibitory receptors, CD96-CD112R, and the immune-stimulating receptor CD226 on immune cells, including CD8^+^ T cells, NK cells and Treg cells. These receptors also competitively or cooperatively interact with several ligands, namely CD155, CD111, CD112, CD113, PVRL4 (25,26). Adding to the complexity, signalling outcomes in immune cells are influenced by the availability of ligands and the kinetics of receptor expression and the binding affinities. Thus, inhibition of tumour expansion by anti-TIGIT Abs is conditioned to the complex network involving TIGIT and its related factors. Notwithstanding, co-expression of CD155 and PD-L1 in non-small cell lung carcinoma (NSCLC) tumours predicts favourable response to PD-1 blockade, as patients with PD-L1 expressing tumours without CD155 were more likely to respond (16). Given the complexity of the TIGIT interactions mentioned above, it is perhaps not surprising that our analysis of expression of PD-L1 and the TIGIT ligand CD155 on multiple PDX samples and cancer cell lines revealed different patterns. For example, analysis of lung cancer PDXs was able to identify PD-L1^lo^CD155^lo^ and PD-L1^hi^CD155^hi^ cancers. It would be important to identify whether these two different types of cancers will respond to combination therapy differently, and whether PD-L1^hi^CD155^hi^ cancers would show favorable response to BsAb (or combination) compared to treatment with anti-PD-L1 Ab alone.

In summary, we demonstrated that BsAb HB0036 displays certain contextual advantages over mAb combination therapy. These advantages include stronger enhancement of T cell proliferation associated with CD226 upregulation, great enrichment of TIGIT Ab at the sites of PD-L1^+^ tumours and increased inhibition of tumour expansion associated with certain favourable immunological characters. Like treatment with anti-PD-1/PD-L1(16), distinguishing which patient cohort is likely to benefit based on the mode of action could be a key to the realization of the therapeutic potential of dual blockade of PD-L1 and TIGIT. Currently, there are several bispecific antibodies targeting PD-1/PD-L1 and TIGIT, including the one described in this study, that have entered clinical trials (27). The findings from this study might help to realize the potential of these bispecific antibodies for the treatment of cancer patients.

## Material and Methods

### Use of mice and human PBMC cells

Mouse experiments were carried out with the approvals of Institutional Animal Care and Use Committee of Gempharmatech Co., Ltd (GPTAP20220413-2; GPTAP20220624-4, GPTAP20220818-1, GPTAP20201214-3, GPTAP011), Biocytogen Pharmaceutical (Beijing) Co. (BAP-BJ-PS-01-2204158), and Pharmalegacy Co. (PL220119-3). The tumour cell lines and mouse strains used in this study were listed in supplemental materials. All the operations and techniques used on these laboratory animals conformed with the rules of laboratory animal welfare and ethics and the "3R principles". All the laboratory animals received humanitarian care throughout the whole research process. For use of the cynomolgus monkey in PK and ADA study, animal care was compliant with the guide of JOINN LABORATORIES (Suzhou) Inc that had been fully accredited by the Association for Assessment and Accreditation of Laboratory Animal Care International (AAALAC). During the test, the animals had been approved by the Science and Technology Department of Jiangsu Province, China. For human PBMCs, ethical were approved by Ethics Committee of Shanghai Zha Xin Integrated Traditional Chinese and Western Medicine Hospital (Approval Scheme No: SL-WBC-202001).

### Generation of monoclonal Abs and bispecific Ab

Characterization of humanized anti-PD-L1 monoclonal Ab (WO2020199860A1, NCT04678908) and humanized anti-TIGIT antibody HB0030 (WO2021227940A1, CTR20212828) had been reported previously. Briefly, the murine antibody variable region genes were cloned from monoclonal hybridoma cells secreting antibodies targeting human PD-L1 ECD or human TIGIT extracellular domain (ECD). The murine antibody genes were then humanized by grafting CDR into human homologous framework. Finally, the humanized antibody variable genes were cloned into expression vectors containing constant regions of human IgG1 heavy chain and kappa light chain. For the preparation of HB0030-ADCC silent antibody, the leucines at 234 and 235 (EU numbering) in the CH2 region of the HB0030 Fc domain were mutated into two alanines (called L234A/L235A) to eliminate antibody’s binding affinity to FcγR and C1 to abolish antibody-dependent cell-mediated cytotoxicity (ADCC) and complement dependent cytotoxicity (CDC) activities. Plasmids containing the corresponding antibody DNA were transfected into CHO-S or CHO-K1 cells. Antibodies from culture supernatants were purified by protein A affinity chromatography.

For generation of bispecific antibody, the variable region sequences of parent anti-PD-L1 and anti-TIGIT (HB0030) were generated by using hybridoma technology. Humanized anti-TIGIT was chosen as the scFv component of the IgG-scFv, the scFv is in VL→ VH orientation, with a (GGGGS)n (n=3,4,5) linker connecting the VL to the VH. A disulphate bond was engineering into anti-TIGIT scFv between heavy chain variable domain VH44 and light chain variable domain VL100 to increase the stability of scFV modules. Anti-TIGIT scFv was linked to the C-terminus of the heavy chain of anti-PD-L1 IgG1 Fc using a flexible peptide linker (G4S)4. The DNA sequences were synthesized by Genewiz company and subcloned into a pcDNA3.1 derived expression vector and amplified in DH5α competent cells. Purified plasmids were then transfected into CHO-K1 cells to develop stable cell lines through the cell line development process using the PEI technique, linearized heavy and light plasmids (50 μg in total, HC: LC=1:2) were used. For the purification of BsAb proteins, the cell lines were cultured in a shaker for seed expansion until the viable cell density of cell lines achieved up to 0.3×10^6^ cells/mL, followed by inoculation in the bioreactors for about 15 days in a fed-batch mode. The antibodies were captured from the cell supernatants using protein A resin (MabSelect SuRe, Cytiva), then purified by anion exchange chromatography (Capto Adhere, Cytiva) and cation exchange chromatography (Poros 50HS, Thermo Fisher).

### Hydrogen-deuterium exchange (HDX) detected by mass spectrometry (MS)

Purified antigen (5 mM), antibody (5 mM) or complex of antigen and antibody (molar ration 1:1) were incubated in buffer (50 mM HEPES, pH7.5, 50 mM NaCl, 5% glycerol, 4 mM MgCl_2_, 2 mM DTT) for 1 hour before the HDX reactions at 4LJ. Samples in 4 μl were then added into 16 μl D2O on exchange buffer (50 mM HEPES, pH 7.5, 50 mM NaCl, 2 mM DTT) and incubated for various time points (e.g., 0,10, 60,300,900s) at 4LJ, then quenched by mixing with 20 μl of ice-cold 3M gHCL, 1% trifluoroacetic acid. Each quenched sample was immediately injected into the LEAP Pal 3.0 HDX platform. Upon injection, samples were passed through an immobilized pepsin column (2mm X 2cm) at 120 ul/min, and the digested peptides were captured on a C18 PepMap300 trap column (Thermo Fisher Scientific) and desalted. Peptides were separated across a 2.1mm X 5cm C_18_ separating column (1.9 um Hypersil Gold, Thermo Fisher Scientific) with a linear gradient of 4% - 40% CH_3_CN and 0.3% formic acid, over 6 min. Sample handling, protein digestion and peptide separation were conducted at 4LJ. Mass spectrometric data were acquired using a Fusion Orbitrap mass spectrometer (Thermo Fisher Scientific) with a measured resolving power of 65,000 at m/z 400. HDX analyses were performed in triplicate, with single preparations of each complex of antigen and antibody. The intensity weighted mean m/z centroid value of each peptide envelope was calculated and subsequently converted into a percentage of deuterium incorporation. Statistical significance for the differential HDX data is determined by an unpaired t test for each time point, a procedure that is integrated into the HDX Workbench software. Corrections for back-exchange were made based on an estimated 70% deuterium recovery, and accounting for the known 80% deuterium content of the deuterium exchange buffer.

The HDX data from all overlapping peptides were consolidated to individual amino acid values using a residue averaging approach. Briefly, for each residue, the deuterium incorporation values and peptide lengths from all overlapping peptides were assembled. A weighting function was applied in which shorter peptides were weighted more heavily and longer peptides were weighted less. Each of the weighted deuterium incorporation values were then averaged to produce a single value for each amino acid. The initial two residues of each peptide, as well as prolines, were omitted from the calculations.

### Surface Plasma Resonance (SPR) Binding Assay

SPR studies were performed on the Biacore 8K™ System (Cytiva, Sweden). The temperature of sample compartment and the flow cell of Biacore 8K system were set to 25°C, and the data collection rate was 10 Hz. The antibodies were captured on sensor chip Protein A (29127556, Cytiva) as ligand. Antibodies were flowed over the sensor chip 60 s at the rate of 10 mL/min. HB0036 was diluted with running buffer to a suitable concentration that could reach the capture level required. To measure the binding affinity to human TIGIT protein, the system was firstly injected with 200nM human PD-L1 (PD1-H5229, ACRO Biosystems) to saturate PD-L1 binding by HB0036, then 20nM human TIGIT (TIT-H52H5, ACRO Biosystems) in 3-fold titrations. To measure the binding affinity to PD-L1, the system was firstly injected with 50nM human TIGIT protein to saturate TIGIT binding by HB0036, followed by the addition of 50nM human PD-L1 in 3-fold titrations. Each analyst (PDL1/TIGIT) was injected for 120 s and dissociated with 600 s at the rate of 30 mL/min. The sensor chip surface was regenerated with 10mM glycine at pH1.5 through a 30 s injection at the rate of 60 mL/min. The binding affinities of the parental antibodies HB0030, αPD-L1 WT IgG1 (900458) and control antibodies (900542 & Atezolizumab) were determined by above mentioned method without any enhancement step. The data were analyzed in the single-cycle kinetic using the capture analysis method, which is pre-defined in the Biacore Insight Evaluation Software. The original data were evaluated in the 1:1 binding model with a fit local kinetics model.

To measure the valence, HB0036, HB0030, and 900458 were diluted with the running buffer to a suitable concentration that could reach the capture level (360 RU for HB0036, 250 RU for HB0030, and 250 RU for 900458). Human PD-L1 proteins and human TIGIT proteins were diluted to a nominal concentration of 200 nM and 50 nM, respectively. The human PD-L1 solution was injected once at the rate of 10 μL/min for 120 s, and then the human TIGIT solution was injected once for 120 s, after the injections the chip surface was regenerated via 10mM glycine at pH1.5. The calculations of the binding stoichiometry of HB0036 and PD-L1 and HB0036 and TIGIT were fit to the following equations:

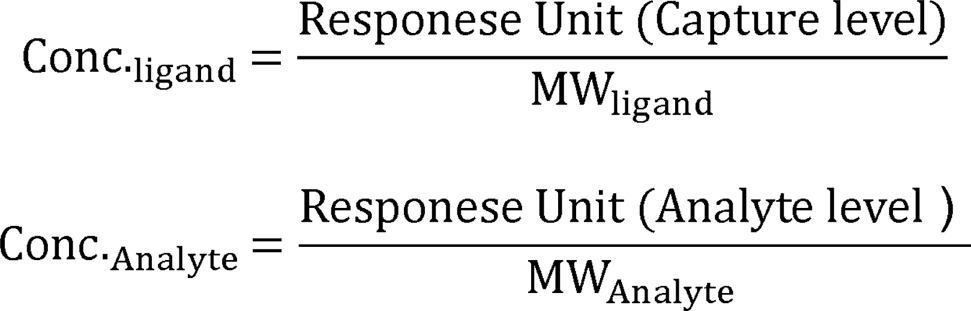

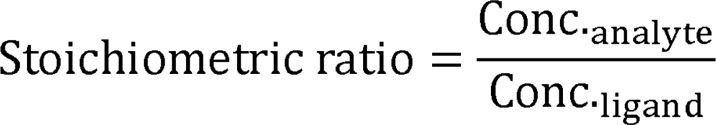

Where, MW_ligand_ and MW_analyte_ are the respective molecular weight (MW) of the ligand and analyte in the running buffer (pH 7.4). The homodimmer structure of PD-L1 or TIGIT has been HPLC-verified by the manufacturer ACRO Biosystems. Therefore, the MW of PD-L1 was determined to be 26000 Da and that of TIGIT was 28800 Da.

### Luciferase Reporter Gene System Assays

Jurkat-NFAT-PD-1-5B8 cells were engineered Jurkat cells that stably express human PD-1 and luciferase reporter under the control of an NFAT promoter. CHO-K1-OS8-PD-L1-8D6 cells were engineered CHO-K1 cells that stably co-express PD-L1 and T-cell activator OKT3 scFv thay could activate Jurkat cells for NFAT transcription and the luciferase expression. Jurkat-NFAT-PD-1-5B8 cells and CHO-K1-OS8-PD-L1-8D6 cells were cultured at 5 × 10^5^/mL in a 96-well plate in 30 μL/well. Serially diluted antibodies were then added into the plates in 30 μL/well. Cells were then incubated at the condition of 5% CO_2_ at 37°C for 6 hrs. After the incubation, the Bio-Glo^TM^ luciferase assay system was applied to estimate the T-cell activation according to the manual (Promega, Madison, WI). The relative luminescence unit (RLU) was detected by using the MD SpectraMax® i3x Microplate Reader scanning at the luminescence unit measure (LUM) endpoint model. Four-parameter fitting was performed with the RLU vs. antibody concentrations using the GraphPad Prism software, and the dose-effect relationship and EC_50_ were determined.

### IL-2 production assay

Jurkat-TIGIT-22G8 cells were engineered Jurkat cells that stably express human TIGIT. Jurkat-TIGIT-22G8 cells produced IL-2 after the stimulation of PHA-M. Jurkat-TIGIT-22G8 cells were cultured 4 × 10^6^/mL in a 96-well plate in 50 μL/well. PHA-M (Yeasen, Cat#40110ES08, lot: P2107890) at final concentration of 0.95 μg/mL were used to activate cells. Soluble CD155 protein (Sino Biological, Cat#10109-H02H) at final concentration of 8 μg/mL were used. Serially diluted antibodies were then added into the plates at the concentration of 50 μL/well. Plates were incubated at 5% CO_2_, 37°C for 22 h. IL-2 secretion was evaluated with the IL-2 assay ELISA kit (Cat#431801, Biolegend). The absorbance value at 450 nm/570 nm was detected by the microplate reader. Four-parameter fitting was performed with the OD_450_-OD_570_ vs. antibody working concentration using GraphPad Prism software, and the dose-effect relationship and EC50 were determined.

### Mixed lymphocyte reaction assay

The CD14^+^ monocytes isolated from PBMCs were cultured with 80 ng/mL GM-CSF (ACRO, Cat#GMF-H4214) and 50 ng/mL IL-4 (R&D, Cat#204-IL-020) for 5 days to generate moDCs. The MLR was performed by seeding 1×10^5^ human CD4^+^ T cells from a different donor 1×10^4^ mo-DC cells in 150 μL/well in the 96-well U-bottom plates. Diluted antibodies were added into the plate at in 50 μL/well. The plates were incubated at 37 °C, 5% CO_2_ for 7 days. Harvested supernatants were assayed for IFN-γ using ELISA kit (Cat#430104, Biolegend).

### In vitro ADCC assay

Human NK cells were seeded at a density of 2×10^6^ cells/mL in a 24-well plate (NUNC) in the presence of IL-15 (10 ng/mL, R&D/247-ILB-025) for overnight. Target cells were CTV-labelled Jurkat expressing huTIGIT. For the assay, target cells (10^5^/well) were plated in 50 μL in a 96-well plate, serially diluted HB0036, HB0030 or isotypes in 50 μL were added into the plate, washed NK cells (10^5^/well) in 50 μL were then added. After incubation for 5 hrs, cells were collected and stained with BV510 anti human CD56 (Biolegend/318340), APC/CY7 anti human CD16 (Biolegend/302017), FITC anti human CD107a antibodies (Biolegend/328606) on ice for 30 min and then subjected to FACS analysis (BD/Canto II) in the presence of Propidium iodide (Sigma/P4170). The rate of cell lysis (%cell lysis) and integrated MFI (iMFI) was respectively calculated by:

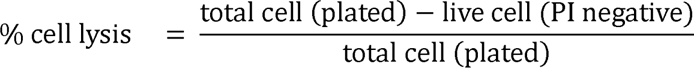

where specifically, %cell lysis was calculated for Jurkat.

### Evaluation of tumour inhibition in vivo

There were two different models used in this study: 1) implantations of human cancer cells in PBMC humanized mice. Human pancreatic adenocarcinoma cells BXPC-3 (10^7^) and equal volume of Matrigel were mixed and inoculated subcutaneously into NPG mice on the right side of the back near the armpit at a volume of about 200 µL. When tumours grew to an average of 50-80 mm^3^, tumour-bearing mice were randomly divided into 4 groups according to tumour volume and body weight, with 6 mice in each group receiving 1×10^7^ PBMC (100uL) via tail vein. Treatment started on the same day. Antibodies were dosed twice weekly for 4 weeks intraperitoneally. 2) implantations of mouse cancer cells in target humanized mice. For evaluation of tumour inhibition by HB0030, with or without anti-PD-1 Ab, CT26 were inoculated subcutaneously on the back of BALB/c-hPD-1/hTIGIT mice. Treatment started when tumours grew to an average of 60-80 mm^3^. For evaluation of tumour inhibition by anti-PD-L1 Ab 900458, 5×10^5^ CT26-hPD-L1 cells were inoculated subcutaneously on the back of BALB/c-hTIGIT/hPD-L1/hPD-1 mice, treatment started when tumours grew to an average of 70 mm^3^. Two additional models were used. for evaluation of tumour inhibition by combination of HB0030 with anti-PD-L1 Ab or bispecific Ab HB0036. One was to inoculate subcutaneously 5×10^5^ MC38-hPD-L1 cells onto the back of hTIGIT/hPD-L1/hPD-1 humanized C57BL/6 mice. Tumour volumes (TV) and body weights were measured and recorded. At the end of experiment, tumour weight was measured and recorded. Analysis of intratumoural infiltrated leukocytes was performed in some experiments.

### Antibodies accumulation at tumour sites in MC38 tumour model

hTIGIT/hPD-L1/hPD-1 humanized C57BL/6 mice were inoculated *s.c*. with 1×10^6^ wild type hPD-L1^-^ MC-38 on the left back near armpit, and 1×10^6^ hPD-L1^+^ MC-38 on the right for 13 days allowing the size of tumour reaching ∼200 mm^3^. Mice were dosed *i.p*. with either 683 μg biotinylated bispicific HB0036 (equal molar concentration to other groups), 500 μg anti-TIGIT HB0030, 500 μg anti-PD-L1 900458, combination of 500 μg HB0030 and 500 μg 900458, 500 μg Control Ig 900201 or PBS. Tumours were collected 24h post Ab injection. Tumour cells and immune infiltrates from tumour digests were then evaluated for antibody accumulation and phenotypes. For FACS based assay, cells were incubated on ice with fixable zombie violet dye (Biolegend/423114), His-tagged TIGIT (ACRO/TIT-H52H5). Cells were then incubated with PE-Streptavidin (Biolegend/405204), BV510 Anti-mouse CD45 (Biolegend/103138), APC/CY7 Anti-mouse CD4 (BD/565650), AF488 anti-mouse FoxP3 (Biolegend/320012), PE/CY7 anti-human TIGIT (Biolegend/372714). Binding of anti-TIGIT antibodies HB0036/HB0030 to cell surface was detected by APC-anti his (Biolegend/362605). For ELISA based assay of anti-TIGIT antibodies in tumour digest, protein TIGIT-his was coated in 96-well plates at 4LJ overnight. Tumour digests were incubated with 2% Tween-20 (Sangon/A100777) to extract proteins. The extract was transferred into TIGIT-his coated plates and incubated for 2 hours, then washed 3 times before incubating with streptavidin-HRP (CST/3999S), then washed 6 times followed by incubating with TMB substrate (Biolegend/421101) and stop solution. Data were expressed by OD_450_.

### Deletion of PD-L1 and CD155 of tumour cells

sgRNAs (PD-L1: 5’TCTTTATATTCATGACCTAC; CD155: 5’CCCGAGCCATGGCCGCCGCG) were chemically synthesized (GenScript). Ribonucleoproteins (RNPs) were produced by complexing sgRNA and recombinant spCas9 (Sino Biological) for 10 min at room temperature. HT1080 cells were mixed with RNPs and subjected to electroporation immediately after complex formation. Harvested cells were then stained for surface PD-L1 and CD155. Four cell populations (PD-L1^+^, CD155^+^, PD-L1^+^CD155^+^, PD-L1^-^CD155^-^) were sorted by flow cytometer and cultured for in vitro study.

### In vitro evaluation of T-cell stimulation with anti-PD-L1 and TIGIT Abs

HT1080, and CD155 KO HT1080, PD-L1 KO, dual-KO HT1080 was harvested and incubated with mitomycin (10 μg/mL) at 37LJ for 30 min. Indicated HT1080 was seeded in 96-well U bottom plate at 2×10^4^/well after washing and incubated with fresh media overnight. Plates after wash were then incubated with antibodies HB0036, Combo (HB0030+900458), HB0030, 900458, 900201. Human PBMCs labelled with CTV was then added at 10^5^/well in media with cocktail of anti-CD3, anti-CD28 antibodies and IL-7. After 4-5 days’ incubation, the cells were then subjected to FACS analysis for activation of T and NK cells while the culture supernatants were evaluated for cytokines. The expression of CD226, CD96, TIGIT, PD-1 in this system was evaluated. In the system without HT1080, soluble CD155 and PD-L1 was incubated with PBMCs. In some cultures, ADCC-silent anti-TIGIT/PD-L1 antibody 700001 and anti-TIGIT antibody were also included.

### Analysis of Tumour infiltrated leukocytes

Tumours were grinded and digested with gentle MACS Octo Dissociator in the presence of DNase LJ. Harvested tumour digests were filtered through 70 μm before centrifuging at 800g for 10 min for cell pellet and resuspended in volumes adjusted according tumour mass. Then cell number were counted. Leukocytes of human and mouse origin were analysed. For spontaneous cytokine release, tumour digests (2×10^5^ cells) were cultured in 96-well plate for 24-hour. Cytokines were evaluated by either ELSIA or CBA based assay.

### Analysis of expression PD-L1, CD155 and related molecules

Data on mRNA expression of PDL1, CD155 and other molecules by PDX and cell lines were downloaded from CROWN BIOSCIENCE (https://www.crownbio.com/databases) and CCLE (https://depmap.org/portal/). Data were classified based on tumour type. Scatter plots with mean and SD were analyzed by GraphPad Prism8.0.1, and each dot indicated data from one PDX or cell line. Correlation between PDL1 and CD155 of several types of tumour were plotted by the same soft. For surface expression by cancer cell lines, cells were stained with indicated antibody. Gated live cells were shown for expression of PD-L and CD155.

### Evaluation of pharmacokinetics (PK) in Cynomolgus monkeys

For single infusion, cynomolgus monkeys (age of 2.6 to 4.5 years, weight around 2.00 to 3.90 kg at the start of experiment) were randomly grouped according to the sex and weight, 3/sex/group. Animals received a single intravenous infusion of 3 mg/kg and 10 mg/kg HB0036 or HB0030, respectively. The duration of infusion was about 20 min. PK: Blood samples from all dose groups were collected from each animal at pre-dose, immediately after dosing finish (± 1 min), 2 h, 6 h, 24 h (D2), 48 h (D3), 72 h (D4), 96 h (D5), 120 h (D6), 168 h (D8), 240 h (D11), 336 h (D15), 408 h (D18), 504 h (D22), 672 h (D29) after dosing. For multiple infusions, cynomolgus monkeys (5/sex/group) were administered saline, or with 10 mg/kg, 30 mg/kg, 100 mg/kg of HB0036 or HB0030, respectively, via intravenous infusion once a week for 4 consecutive weeks (total 5 doses). The blood samples were collected from animals in saline group at pre-dose and 2 hours after the beginning of infusion on Day 1 and Day 22, and treated at pre-dose, immediate completion of infusion (±1 min), 2, 6, 24, 48, 72, 120 and 168 hours after the beginning of infusion on Day 1 and Day 22. In addition, the blood samples were collected from all animals of treated groups at pre-dose on Day 15 and 2 hours after the beginning of infusion on Day 8, Day 15 and Day 29. The concentrations of HB0036 or HB0030 in serum were determined using a validated ELISA method (Study Number: P21-S078-PKV) with a lower limit of quantification of 0.00500 μg/mL. Pharmacokinetic parameters of HB0036 in each animal were calculated using non-compartmental analysis (NCA) in WinNonlin 8.0.0.3176.

### Statistical Analyses

The results are presented as mean ±standard error of the mean (SEM). p < 0.05 was considered to indicate statistical significance (* p < 0.05, ** p < 0 .01, and *** p < 0 .001 using the PRISM 9 one-way ANOVA, followed by the least significant difference multiple comparison test). The differences in the TVs between the compared groups were analyzed by two-way ANOVA.

## Acknowledgement

This work was supported by grants from Shanghai Rising-Star Program (No: 23QB1400400 to JG Xu). We thank Drs Andrew M Lew, Andreas Strasser, Michael Chopin, Michael Zhan and Jenny Zhan for comments and proof-reading the manuscript.

## Supplemental Figures

**Figure S1 (related to Figure 3)**. (A) Schematic illustration of the assay setup. FACS plots (bottom) show expression of PD-L1 and CD155 on WT HT 1080, PD-L1/CD155 double KO, PD-L1 KO and CD155 KO clones. (B) Line graphs show recovered numbers of proliferating cells in the presence of serially diluted antibodies. Histograms show cell division with serially diluted HB0036 or presence of HB0030 and 900458. (C) Bar graphs show number of proliferated cells from cultures with various variants of HT1080 cells. (D) Bar graphs show numbers of proliferated cells from cultures with soluble PD-L1 and CD155. (E). Representative FACS plots show expression of CD226 on gated CD4^+^ and CD8^+^T cells.

**Figure S2 (related to Figure 3)**. Data were set up as in Figure 3E. (A) Representative FACS plots show the proportions of IFN-γ^+^ cells within gated CD4^+^ T cells, CD8^+^ T cells or CD56^+^ NK cells. (B) Bar graphs show summarized data of proportions of IFN-γ^+^ cells under different culture conditions. **P<0.01 by One-way ANOVA multiple parameter comparison. (C). Representative FACS plots show CD107a expression by NK cells in the presence of different Abs.

**Figure S3 (related to Figure 4). Experiment set up as for Figure 4**. hTIGIT/hPD-L1/hPD-1 humanized mice were inoculated with hPD-L1^+^ and hPD-L1^-^ WT MC38 cells on the contralateral side, kept for 13 days and theen injected with the indicated biotinylated Abs for 1 day. mCD45^+^ cells from tumour digests were analyzed for CD4^+^ T cells. A. Evaluation of intra-tumoural CD4^+^ T cells amongst the mCD45^+^ cells from tumour digests. Bar graphs show mean percentages+/-SEM of CD4^+^ cells. B. CD4^+^ T cells from intra-tumoural mCD45^+^ cells were analysed for Foxp3 expression. FACS plots show Foxp3^+^ Tregs within CD4^+^ cells. Bar graphs show % (Mean+/-SEM) of Tregs among CD4^+^ cells. ** P<0.01, ns (not significant) by One-way ANOVA multiple parameter comparison. D. Detection of anti-TIGIT Ab binding to TIGIT by CD4^+^ cells. TIGIT-His was used to detect potential binding of injected anti-TIGIT Ab by above CD4^+^ T cells.

**Figure S4 (related to Figure 5&6). Inhibition of tumor eexpansion by bi-specific mAbs and mAb combination**. Tumour cells were inoculated into the back of mice. Treatment started when tumour volumes had reached the indicated size. Abs (mg/kg body weight) were administered by intraperitoneal injection twice weekly for total 5 doses. Data of tumour volumes and tumour weights are presented. In some experiments, data of intra-tumoural leukocytes are provided. *P<0.05, **P<0.01 by One-way Anova multiple parameter comparison. (A) Inhibition of tumour expansion by anti-TIGIT mAb HB0030. BALB/c-hPD1/hTIGIT mice were inoculated with 5×10^5^ CT26 tumour cells. The treatment was started when the tumour size had reached an average TV of 60 mm^3^. The treatment groups included vehicle control, HB0030 or HB0030 ADCC silent antibody at 3 mg/kg body weight (n=6). (B) Inhibition of tumour growth by anti-PD-L1 mAb 900458. hPD-L1/hPD-1 humanized BALB/c mice were inoculated s.c. with 5×10^5^ CT26-hPD-L1 cells allowing the size of tumour to reach 70 mm^3^. Control Ig 900201 or anti-PD-L1 900458 at 1 or 3 mg/kg body weight (n=8) were. (C) Inhibition of tumour expansion by combination of HB0030 with Keytruda. BALB/c-hPD-1/hTIGIT mice were inoculated with 5×10^5^ CT26 tumour cells and tumours were allowed to reach an average TV 80 mm^3^. Groups (n=6) included control 900201, HB0030, Keytruda, or combination HB0030 and Keytruda. Doses were 10 mg/kg body weight for all except Keytruda which was 3 mg/kg body weight. (D) Inhibition of tumour growth by combination of HB0030 with anti-PD-L1 Ab 900458. hPD-1/hPD-L1/hTIGIT humanized C57BL/6 mice were inoculated with 5 x 10^5^ MC38-hPD-L1 cells and tumours were allowed to reach an average TV 100 mm^3^, Treatment groups (n=6) included control 900201, HB0030, 900458 or combination with HB0030 and 900458 at 3 mg/kg body weight.

**Figure S5 (related to Figure 5)**. Analysis of TILs in the syngeneic CT26-hPDL1 mouse model with hTIGIT/hPD-L1 humanization BALB/c-hPD1/hPDL1/hTIGIT mice. A. Gating of CD45^+^ TILs. B. Gating of myeloid cell subsets from CD45^+^ TILs. C. Gating of T cell subsets from CD45^+^ TILs. D. Gating of CD226 expression on T cell subsets. E. Gating of CD49^+^ NK cells from CD45^+^ TILs. Bar graph shows relative percentages of NK cells. Linear regression between the relative percentages of NK cells and and tumour weights was also shown. F. Gating of cDC1s and pDCs from CD45^+^ TILs. Bar graphs show relative percentages of cDC1s and pDCs. Linear regression between relative percentages of DC subsets and tumour weights is also shown.

**Figure S6 (related to Figure 5&6) A**. Analysis of TILs harvested from BxPC-3 tumours. FACS plots show gating of expression of CD226 and CD96 on the indicated T cell subsets. **B**. Inhibition of tumour expansion in PBMC humanized mice bearing HT1367 cancers. NPG mice received human bladder carcinoma cells HT1376 (1×10^7^) on the right flank. When tumors had grown to an average size of 50-80 mm^3^, tumor bearing mice were randomly divided into groups according to tumor volume and body weight, with 6 mice in each group receiving 1×10^7^ PBMC. Treatment started on the same day. Antibodies were dosed twice weekly for 4 weeks intraperitoneally. *P<0.05, ***P<0.001, ****P<0.0001 between HB0036 and combination groups by 2-way ANOVA raw comparison. **(C-D)**. Comparison of PK and ADA of HB0036 and HB0030 in mouse. C57BL/6 mice (n=3) were given the indicated concentration of Abs (HB0036 were adjusted to equal molar amounts). Sera were collected for evaluation of drug concentration and ADA. B. Drug concentration, expressed as % of maximal amounts detected (1 day after injection), are shown. C. ADA titers at 21 days after injection are shown. **(E-F)** Comparison of PK of HB0036 and HB0030 in naïve Cynomolgus monkeys. E. Cynomolgus monkeys (n=3 each for both male and female).

**Figure S7 (related to Figure 7)**. *CD155* mRNA expression in cancer PDX and cancer cell lines (https://hubase.crownbio.com/GeneExpressionRNAseq/Index <u>and</u> https://xenobase.crownbio.com/GeneExpressionRNAseq/Index) were analyzed. (A) Co-expression pattern of PD-L1 and CD155 in cell line samples. (B) Expression of the CD155 related molecules CD111, CD112, CD113 and PRVL4 by PDX samples. (C). Expression of the CD155 related molecules CD111, CD112, CD113 and PRVL4 in cancer cell lines.

