## Supplementary figures and images for "Co-engagement of TIGIT^+^ immune cells to PD-L1^+^ tumours by a bispecific antibody potentiates T-cell response and tumour control"

### Supplemental Figure 1

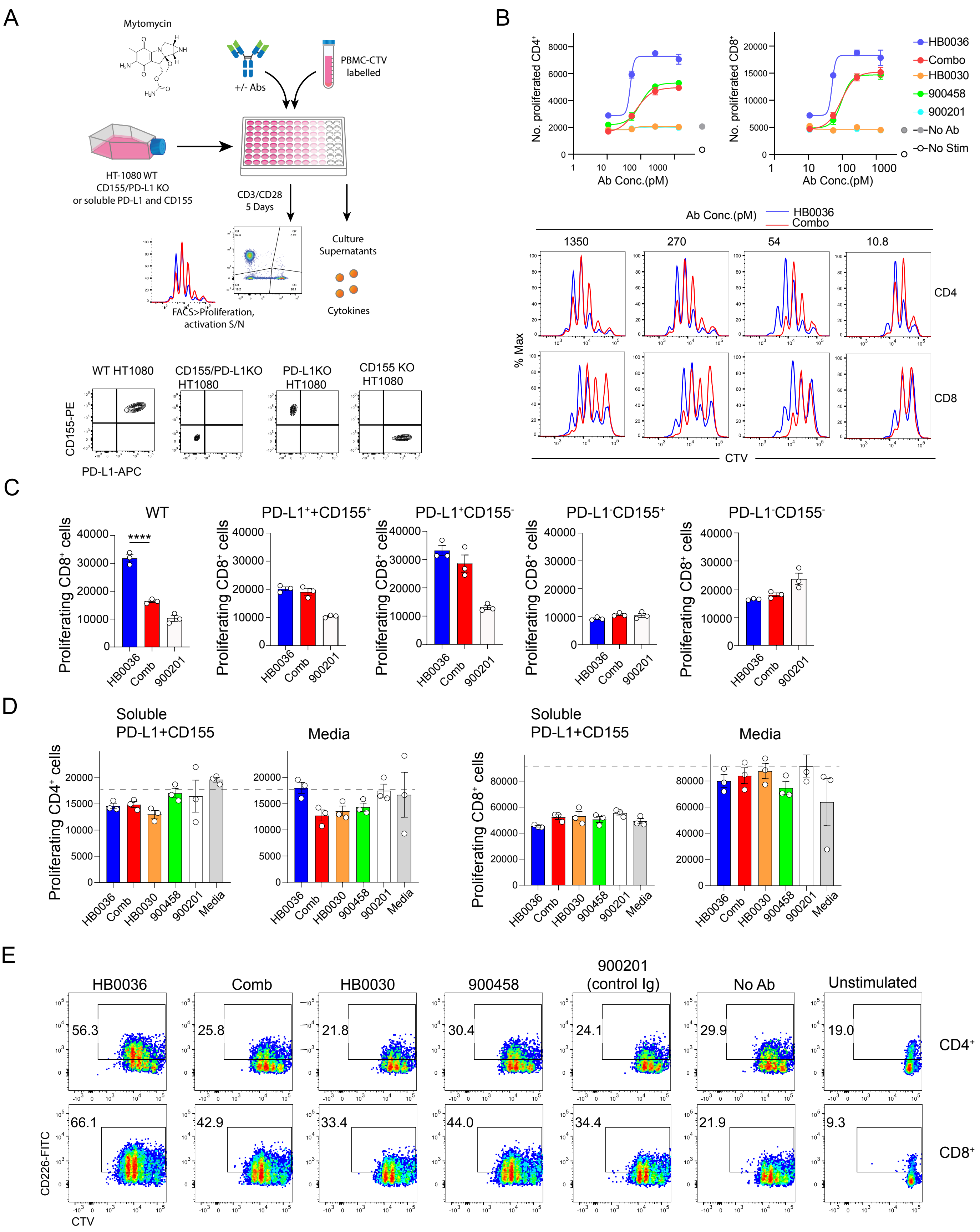

### Supplemental Figure 2

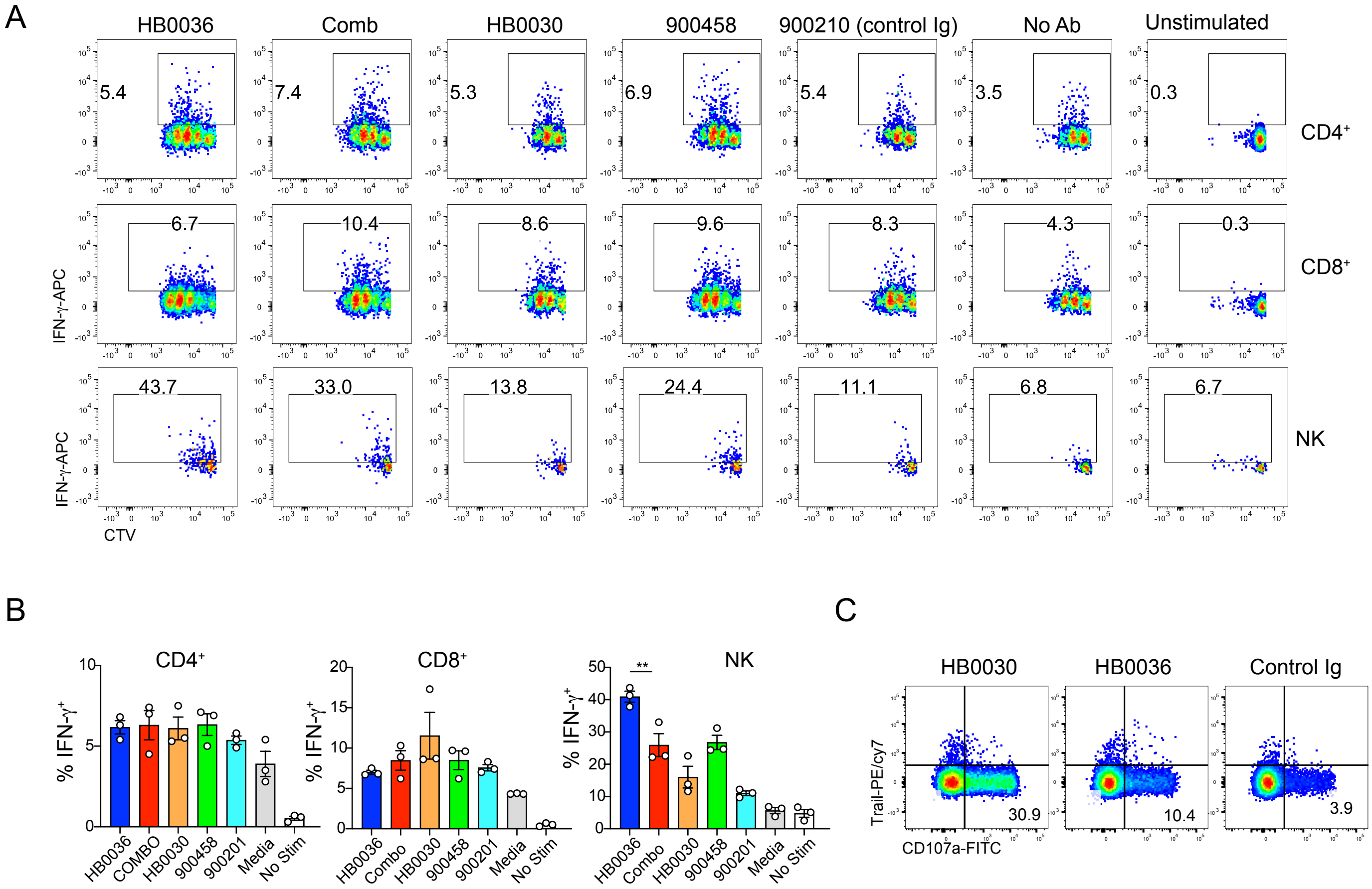

### Supplemental Figure 3

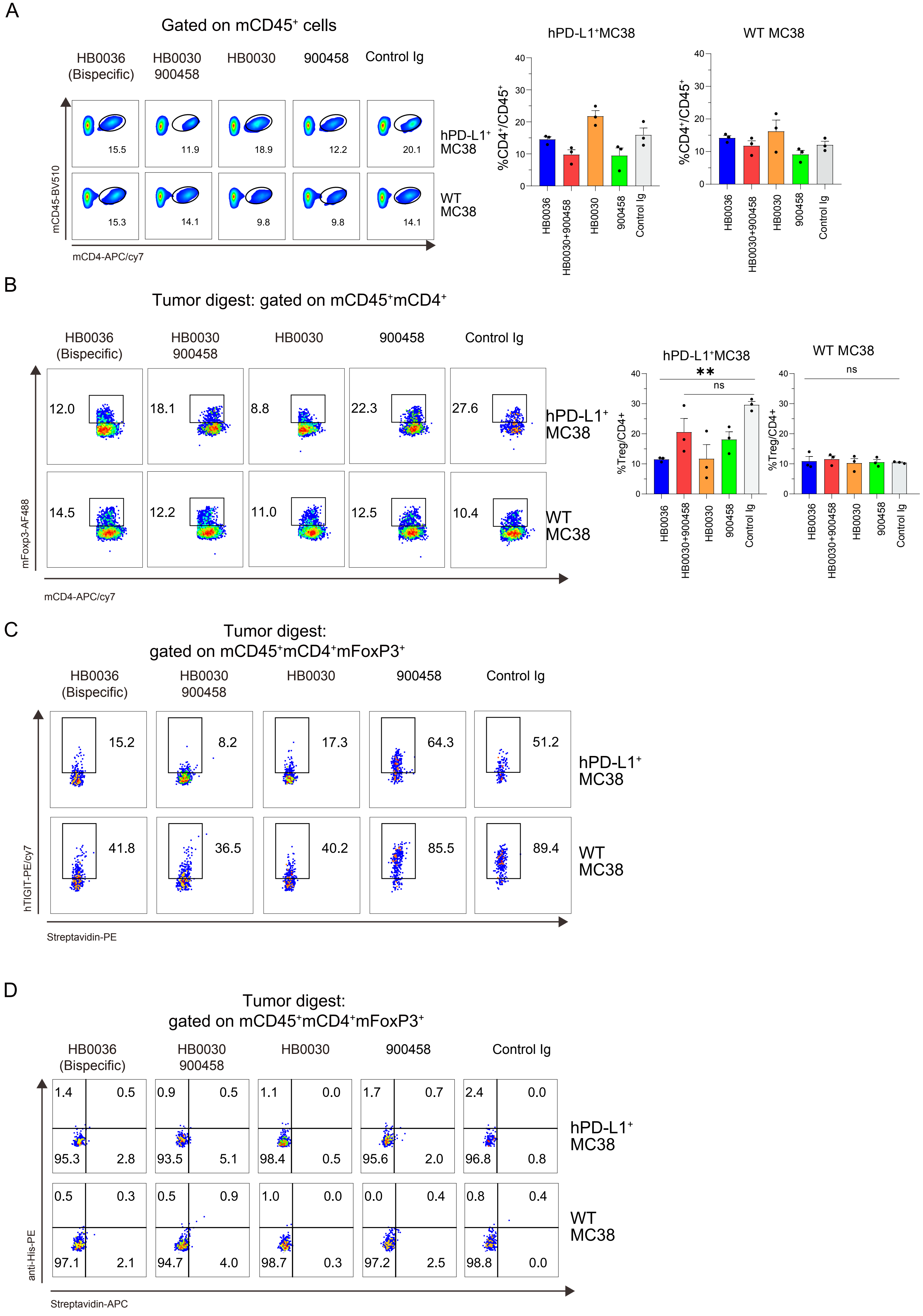

### Supplemental Figure 4

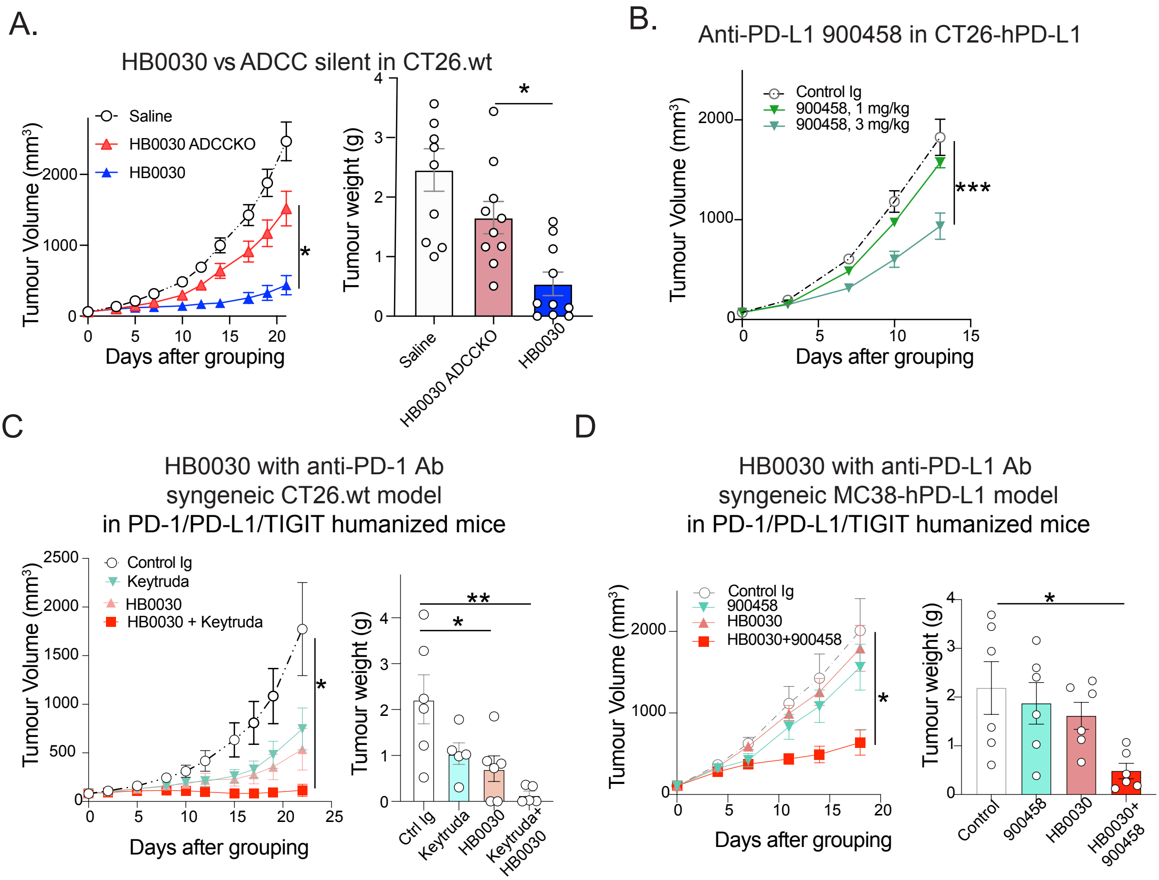

### Supplemental Figure 5

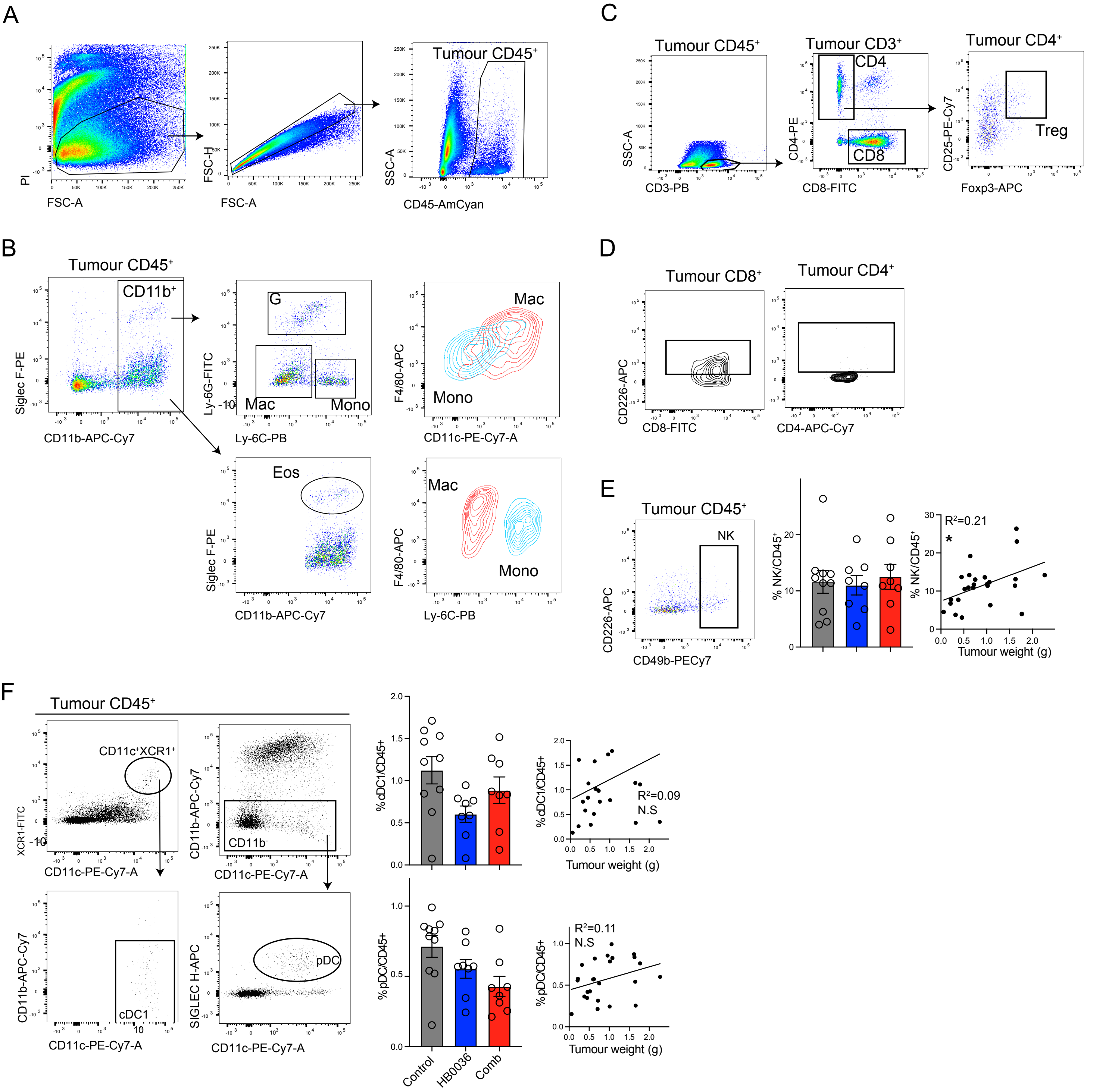

### Supplemental Figure 7

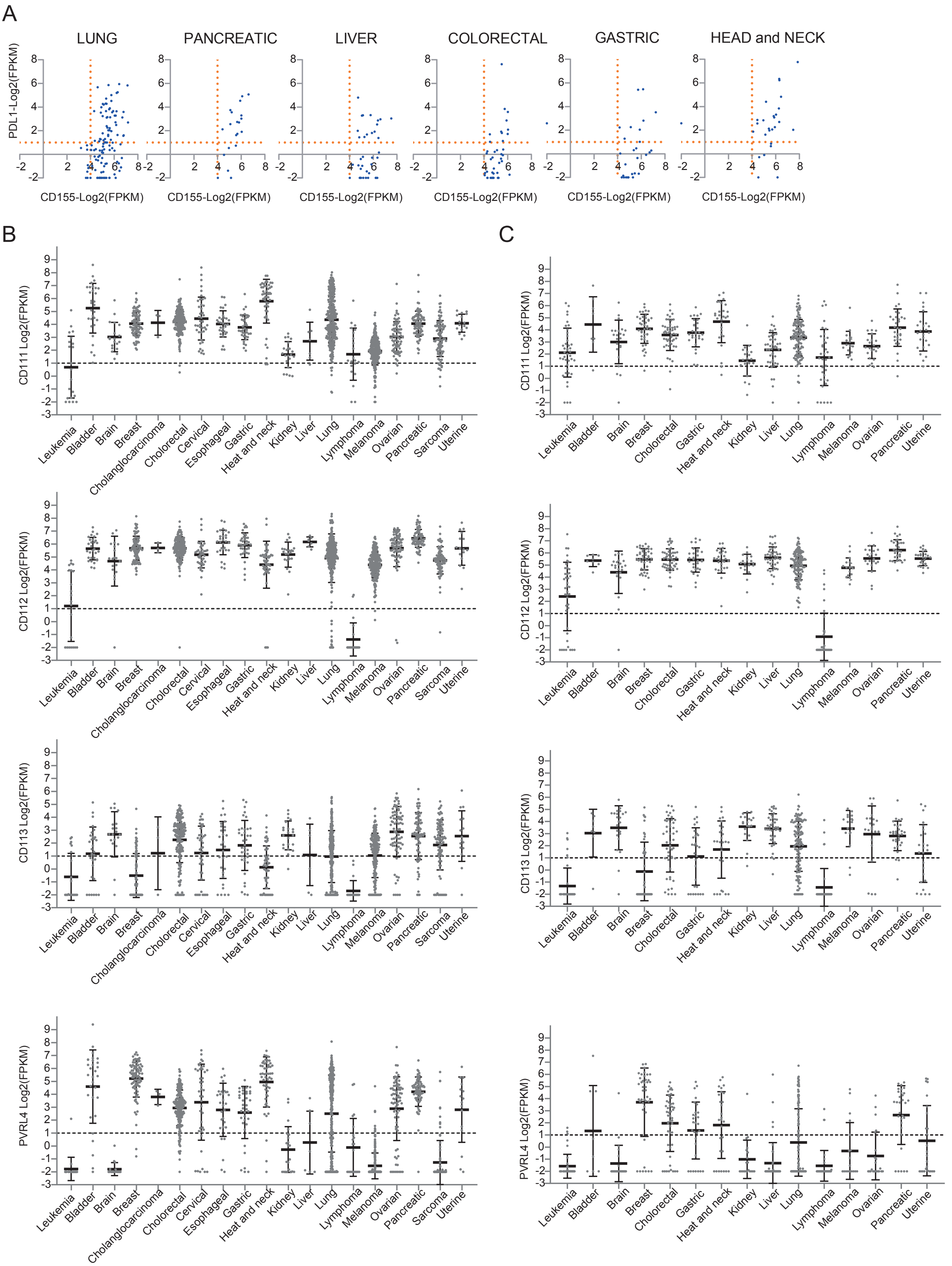
