## Supplemental Figure 6 for "Co-engagement of TIGIT^+^ immune cells to PD-L1^+^ tumours by a bispecific antibody potentiates T-cell response and tumour control"

Figure 6S

A Analysis of TILs

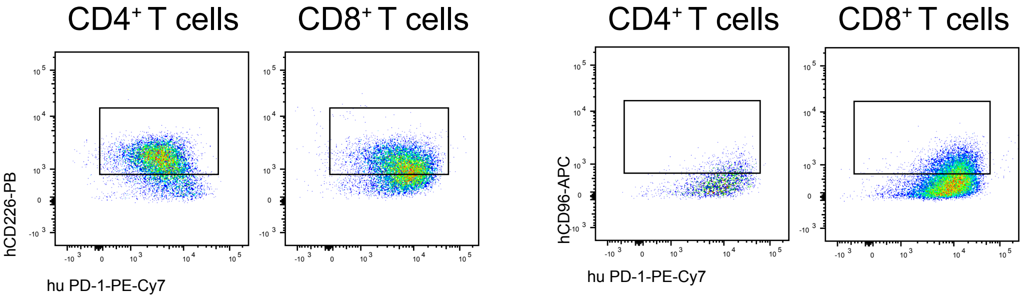


B tumour inhibition was also observed in xenograft with
human bladder carcinoma cell line HT1367 in PBMC humanized mice

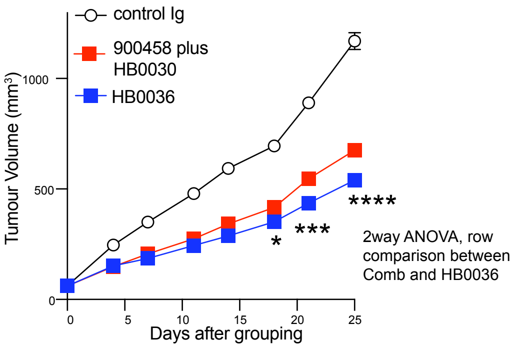


C PK in WT mouse

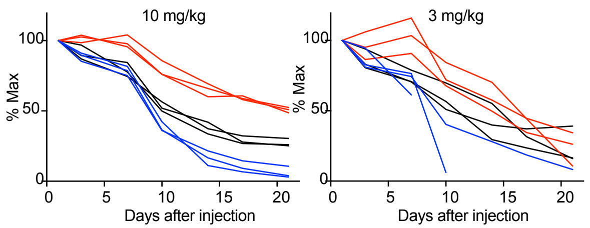


D ADA in WT mouse

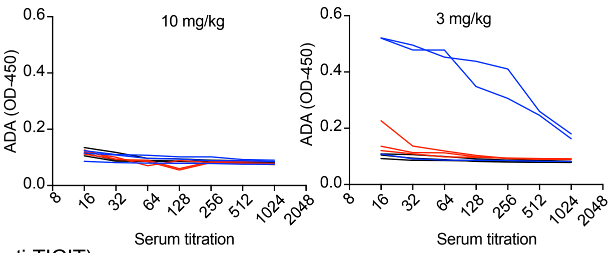


HB0030 (anti-TIGIT)
HB0036 (Bispecific)
900458 (anti-PD-L1)

E. PK after single infusion
in cynomolgus monkeys

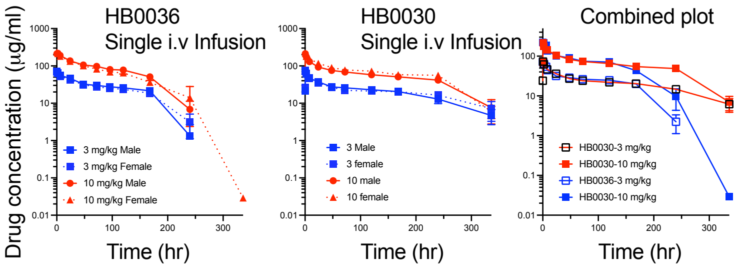


F. PK after multiple infusions
in cynomolgus monkeys

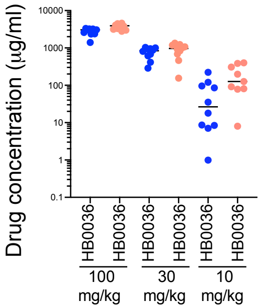


2 hr after last dose at day 29

Animal received dosing
at Day 1, 8, 15, 22, 29
