## Supplemental Table 1 for "Co-engagement of TIGIT^+^ immune cells to PD-L1^+^ tumours by a bispecific antibody potentiates T-cell response and tumour control"

**Table S1** Structural features, SEC purity and thermodynamic properties of PD-L1/TIGIT BsAbs

| PD-L1/TIGIT BsAbs | Length of linker 2 | Disulfide-stabilized scFv | SEC purity（%） | Tm value (℃) | Tagg value (℃) |
| --- | --- | --- | --- | --- | --- |
| 900691 | 15 | VH44-VL100 | 65.82 | 69.2 | 66.5 |
| 900692 | 20 | VH44-VL100 | 85.21 | 69.4 | 67.1 |
| 900693(HB0036) | 25 | VH44-VL100 | 90.41 | 70.1 | 67.5 |
