## Supplemental Table 2 for "Co-engagement of TIGIT^+^ immune cells to PD-L1^+^ tumours by a bispecific antibody potentiates T-cell response and tumour control"

**Table S2: Antibodies, biological materials and resources**

| **REAGENT or RESOURCE** | **SOURCE** | **IDENTIFIER** |
| --- | --- | --- |
| **Antibodies (Clone)** | | |
| HB0036 | In house | N/A |
| HB0030 | In house | N/A |
| HB0030-ADCC KO | In house | N/A |
| HB0030-ADCC enhanced | In house | N/A |
| 900458 | In house | N/A |
| 900201 (Ctl Ig) | In house | N/A |
| 900543 | In house | N/A |
| 700001 | In house | N/A |
| 900541 | In house | N/A |
| 900542 | In house | N/A |
| Atezolizumab | Roche | H0211 |
| Tecentriq (provided by Gempharmatech) | Roche | Lot# H0217B01 |
| Keytruda | MSD | Lot# S001195 |
| APC anti-His Tag (J095G46) | BioLegend | Cat#362605, RRID:AB_2715818 |
| PE anti-His Tag (J095G46) | BioLegend | Cat#362603, RRID:AB_2563634 |
| BV510 anti-mouse CD45 (30-F11) | BioLegend | Cat#103138, RRID:AB_2563061 |
| APC/CY7 Anti-mouse CD4 (RM4-5) | BD | Cat#565650, RRID:AB_2739324 |
| AF488 anti-mouse FoxP3 (150D) | BioLegend | Cat#320012, RRID:AB_439748 |
| PE/CY7 anti-human TIGIT (A15153G) | BioLegend | Cat#372714, RRID:AB_2632929 |
| FITC anti-human CD107a (H4A3) | BioLegend | Cat#328606, RRID:AB_1186036 |
| BV510 anti-human CD56 (HCD56) | BioLegend | Cat#318340, RRID:AB_ 2561944 |
| APC/CY7 Anti-human CD16 (3G8) | BioLegend | Cat# 302017, RRID:AB_ 314217 |
| FITC anti-mouse Ly-6G Antibody(1A8) | BioLegend | Cat#127606, RRID:AB_ 1236494 |
| PE-Cy™7 Hamster Anti-Mouse CD11c(HL3) | BD | Cat#558079, RRID:AB_ 647251 |
| APC Anti-Mouse F4/80 (BM8) | eBioscience | Cat#17-4801-80, RRID:AB_ 2784648 |
| APC-Cy™7 Anti CD11b (M1/70) | BD | Cat#557657, RRID:AB_ 396772 |
| Pacific Blue™ anti-mouse Ly-6C (HK1.4) | BioLegend | Cat#128014, RRID:AB_ 1732079 |
| Alexa Fluor® 488 anti-human CD8 (SK1) | BioLegend | Cat#344716, RRID:AB_ 10549301 |
| PE anti-human CD56 (NCAM) | BioLegend | Cat#318306, RRID:AB_ 604101 |
| PerCP-Cyanine5.5 anti-human CD3 (UCHT1) | Invitrogen | Cat#MA5-38706, RRID:AB_2898618 |
| APC anti-human CD45RO (UCHL1) | BioLegend | Cat# 304210, RRID:AB_ 314426 |
| APC/Cyanine7 anti-human CD4 (OKT4) | BioLegend | Cat# 317418, RRID:AB_ 571947 |
| Brilliant Violet 421™ anti-human CD45 (2D1) | BioLegend | Cat# 368522, RRID:AB_ 2687375 |
| Anti-Hu CD28, eBioscience (CD28.2) | Thermo | Cat# 14-0289-82, RRID:AB_467194 |
| LEAF™ Purified anti-human CD3 Antibody (HIT3a) | Biolegend | Cat# 300332, RRID:AB_11150396 |
| PE anti-human CD8 Antibody (SK1) | Biolegend | Cat# 344706, RRID:AB_1953244 |
| FITC anti-human CD226 (DNAM-1) Antibody (11A8) | Biolegend | Cat# 338304, RRID:AB_2228763 |
| APC anti-human TIGIT(VSTM3)Antibody (A15153G) | BioLegend | Cat# 372705, RRID:AB_2632731 |
| PE/Cyanine7 anti-human CD96 (TACTILE) Antibody (NK92.39) | BioLegend | Cat# 338415, RRID:AB_2629523 |
| Brilliant Violet 510™ anti-human CD8 Antibody | BioLegend | Cat# 344731, RRID: AB_2564623 |
| PE Mouse Anti-Human CD279 | BD Pharmingen | Cat#557946, RRID: AB_647199 |
| FITC anti-mouse/rat XCR1 Antibody | BioLegend | Cat#148210, RRID: AB_2564366 |
| PE anti-mouse CD4 Antibody | BioLegend | Cat#100408, RRID: AB_312693 |
| APC anti-mouse Siglec H Antibody | BioLegend | Cat#129612, RRID: AB_10641134 |
| [BD Pharmingen™ PE Mouse Anti-Human CD274](https://www.bdbiosciences.com/zh-cn/products/reagents/flow-cytometry-reagents/research-reagents/single-color-antibodies-ruo/pe-mouse-anti-human-cd274.561787) | BD Pharmingen | Cat# 561787, RRID: AB_647198 |
| [BD Pharmingen™ PE Rat Anti-Mouse Siglec-F](https://www.bdbiosciences.com/zh-cn/products/reagents/flow-cytometry-reagents/research-reagents/single-color-antibodies-ruo/pe-rat-anti-mouse-siglec-f.552126) | BD Pharmingen | Cat# 552126, RRID: AB_394341 |
| Brilliant Violet 421™ anti-mouse Ly-6G/Ly-6C (Gr-1) Antibody | BioLegend | Cat# 108434, RRID: AB_2562219 |
| BD Horizon™ BV421 Hamster Anti-Mouse TCR β Chain | BD Pharmingen | Cat# 562839, RRID: AB_2737830 |
| FITC anti-mouse CD8a Antibody | BioLegend | Cat# 100706, RRID: AB_312745 |
| PE/Cyanine7 anti-mouse CD49b (pan-NK cells) Antibody | BioLegend | Cat# 108922, RRID: AB_2561460 |
| CD226 (DNAM-1) Monoclonal Antibody (10E5), APC, eBioscience™ | Invitrogen | Cat# 17-2261-82, RRID: AB_11149875 |
| PE anti-mouse CD96 (TACTILE) Antibody | BioLegend | Cat# 131705, RRID: AB_1279389 |
| Pacific Blue™ anti-mouse CD3 Antibody | BioLegend | Cat# 100214, RRID: AB_493645 |
| BD Pharmingen™ PE-Cy™7 Rat Anti-Mouse CD25 | BD Pharmingen | Cat# 552880, RRID: AB_394509 |
| Alexa Fluor® 647 anti-mouse/rat/human FOXP3 Antibody | BioLegend | Cat#320014, RRID: AB_439750 |
| **Bacterial and virus strains** | | |
| DH5α competent cells | Yeastern Biotech | Cat# FYE607-80VL |
|  |  | Cat# |
| **Oligonucleotides** | | |
|  |  | Cat# |
|  |  | Cat# |
| **Biological Samples** | | |
| Human NK cells | TPCS | Cat# PB56-N-1CW |
| Jurkat-FcγRIIIa(158V) | Genomeditech | NA |
| PBMC | Sailybio | SLB-HP010B/200273&200339 |
| hPBMC | TPCS | Cat# PB010C-W |
| **Chemicals, Peptides, and Recombinant Proteins** | | |
| Human TIGIT Protein, His Tag (MALS verified) | ACRO Biosystems | Cat# TIT-H52H5 |
| Human PD-L1 / B7-H1 Protein, His Tag (MALS verified) | ACRO Biosystems | Cat# PD1-H5229 |
| Zombie violet™ Fixable Viability Kit | BioLegend | Cat#423114 |
| Zombie Aqua™ Fixable Viability Kit | BioLegend | Cat#423102 |
| Zombie NIR™ Fixable Viability Kit | BioLegend | Cat# 423106 |
| PE Streptavidin | BioLegend | Cat#405204 |
| APC Streptavidin | BioLegend | Cat#405207 |
| Propidium iodide | Sigma-Aldrich | Cat#P4170 |
| Tween-20 | Sangon | Cat#A100777 |
| streptavidin-HRP | CST | Cat#3999S |
| TMB substrate | BioLegend | Cat#421101 |
| Recombinant Human IL-15 Protein | R&D | Cat#247-ILB-025 |
| HBS-EP+(10×)，pH7.4 | Cytiva | Cat# BR-1006-69 |
| 10mM Glycine，pH1.5 | Cytiva | Cat#BR-1003-54 |
| Series S Sensor Chip Protein A | Cytiva | Cat#29127556 |
| GM-CSF | ACRO | Cat# GMF-H4214 |
| IL-4 | R&D | Cat#204-IL-020 |
| IFN-γ ELISA kit | BioLegend | Cat#430104 |
| MEM | Gibco | Cat# 11095-080 |
| RPMI 1640 | Gibco | Cat# A1049101 |
| Recombinant Human IL-7 Protein | R&D | Cat# 207-IL-025/CF |
| Mitomycin for Injection | HISUN | Cat# H19999025 |
| **Critical Commercial Assays** | | |
| True-Nuclear Transcription Factor Buffer Set | Biolegend | 424401 |
| CellTrace Violet Cell Proliferation Kit | Thermo | C34557 |
| ELISA MAX™ Deluxe Set Human IFN-γ | Biolegend | Cat#430104 |
| LEGENDplex™ Mouse Inflammation Panel (13-plex) with V-bottom Plate | Biolegend | Cat#740446 |
| LEGENDplex™ Human CD8/NK Panel (13-plex) with V-bottom Plate | Biolegend | Cat#741065 |
| Bio-Glo^TM^ Luciferase assay | Promega | G7940 |
| Cytotoxicity LDH Assay Kit-WST | Dojindo | Cat#423114 |
| **Deposited data** | | |
| Data will be made publicly accessible upon publication |  |  |
| **Experimental Models: Cell Lines** | | |
| Jurkat-huTIGIT-22G8 | In house | N/A |
| MC38-hPD-L1 | Pharmalegacy | N/A |
| MC38 | Pharmalegacy | N/A |
| BxPC-3 | Pharmalegacy | N/A |
| CT26-hPD-L1 | Gempharmatech | N/A |
| CT26 | Gempharmatech | N/A |
| EMT6- hPD-L1 | Gempharmatech | N/A |
| MC38-hPD-L1 | Biocytogen | N/A |
| CHO-K1-PD-L1-8C1 | In house | N/A |
| NK-92MI-CD16a | In house | N/A |
| HT-1080 | ATCC | Cat# CCL-121 |
| hPDL1 & hCD155 knockout HT-1080 | 赛斯尔擎 | N/A |
| **Experimental Models: Organisms/Strains** | | |
| Mouse: C57BL/6-hPD-1/hPD-L1/hTIGIT | Pharmalegacy | Cat# 130573 |
| Mouse: NOD.Cg-Prkdc^scid^ Il2rg^tm1Vst^/Vst | Pharmalegacy | Cat# |
| Mouse: BALB/c-hPD-1/hPD-L1 | Gempharmatech | NO. T004025 |
| Mouse:BALB/c-hPD1/hTIGIT | Gempharmatech | Cat# T004023 |
| Mouse:BALB/c-hPD1/hPD-L1/hTIGIT | Gempharmatech | Cat# T037004 |
| Mouse:Balb/C | Pharmalegacy | Cat# |
| **Recombinant DNA** | | |
| DNA coding for anti-PD-L1 | Genewiz | N/A |
| DNA coding for anti-TIGIT scFv | Genewiz | N/A |
| DNA coding for (GGGGS)n (n=3,4,5) linkers | Genewiz | N/A |
| **Software and Algorithms** | | |
| Flowjo V10 | FlowJo, LLC | <https://www.flowjo.com/> |
| Prism 8.0 | GraphPad Software | <https://www.graphpad.com/scientific-software/prism/> |
| Adobe Illustrator CC 2019 | Adobe | https://www.adobe.com/sea/products/illustrator.html?promoid=PGRQQLFS&mv=other |
| LEGENDplex™ | BioLegend | https://legendplex.qognit.com/workflow |
| **Other** | | |
| BD FACSCanto II | BD Bioscience | N/A |
| BD FACSMelody | BD Bioscience | N/A |
| gentleMACS Octo Dissociator with Heaters | Miltenyi Biotec | 130-096-427 |
| Biacore 8K | Cytiva | N/A |
| Microplate reader | Molecular Device | I3x |
